# Reconstructing sequence-grammar trajectories enables interpretable and tunable *cis*-regulatory element design

**DOI:** 10.64898/2026.07.10.737719

**Authors:** Mingqian Ma, Wanjuan Bu, Guoqing Liu, Yuxuan Liu, Sizhen Liu, Zhen Zhao, Shijie Yao, Qingru Hua, Yujie Zhang, Cuiting Zhong, Haitao Huang, Pan Deng, Peiran Jin, Qijin Yin, Chuan Cao, Haiguang Liu, Mo Xu, Yuan He, Tao Qin, Zeyu Chen

## Abstract

Designing synthetic *cis*-regulatory elements (CREs) with cell-type-specific activity remains challenging, and optimization is usually treated as a black box, obscuring how regulatory grammar emerges and why design trajectories fail. Here, we present GO-CRE (<u>G</u>uided <u>O</u>ptimization of *<u>C</u>is*-<u>R</u>egulatory <u>E</u>lements), which combines efficient sequence generation, predictor-guided reinforcement learning, and trajectory-level interpretation. GO-CRE reconstructs iterative sequence changes in a shared sequence-grammar landscape and identifies coordinated update programs corresponding to search, commitment, and optimization. Productive trajectories in HepG2 and K562 progressively acquired cell-type-associated grammar, whereas SK-N-SH trajectories remained confined to local basins. In HepG2, trajectory analysis revealed a low-complexity polyG trap; introducing a polyG penalty redirected optimization toward HNF/FOXA-associated features. Final designs retained sequence diversity while converging on cell-type-associated motif patterns. Lentiviral MPRA validated cell-type-specific activity in K562 and HepG2 and showed higher average activity of HepG2 designs than endogenous CREs. Together, these findings establish sequence-grammar trajectory reconstruction as a basis for interpretable and tunable synthetic CRE design.

## Introduction

*Cis*-regulatory elements (CREs) are central regulators of context-specific gene expression, shaping cell fate decisions, cell state transitions, and cell type specification^1–4^. Genome-wide chromatin landscape profiling has revealed broad associations between CRE states and gene expression^5,6^, whereas genetic and epigenetic perturbation studies have provided causal evidence for how specific CREs control gene expression in a context-dependent manner^7–15^. Recent advances in genomic language models (gLMs) have expanded our ability to decode DNA-encoded regulatory features across organisms and cell types, particularly for understanding CRE function and regulation^16–30^. Many of these models use encoder-based architectures that are well suited to capturing contextual information within DNA sequences, and several highly interpretable models have provided base-resolution insights into regulatory mechanisms. These advances have transformed our ability to interpret endogenous regulatory sequences. However, the inverse problem, the *de novo* design of synthetic CREs for context-specific or user-defined functions, remains substantially less mature.

Synthetic CRE design has important applications in functional genomics and programmable gene control, particularly when endogenous regulatory repertoires are incomplete, insufficiently selective, or poorly suited to therapeutic delivery. Recent approaches have begun to address this challenge through either experimental directed evolution in mammalian cells or *in silico* generative design. *In situ* directed-evolution strategies couple mutagenesis to functional selection in native cellular environments, but they are constrained by limited mutational diversity, locus throughput and scalability^31–33^. *In silico* generative models, including convolutional neural networks, diffusion models and autoregressive genomic language models, can evaluate substantially larger candidate pools and have generated synthetic DNA sequences with cell-type-specific regulatory activity^24,34–37^. More recently, iterative deep-learning design has coupled model-guided sequence generation with experimental testing and model refinement, producing a second generation of human enhancers with improved cell-type specificity and a more condensed motif vocabulary than endogenous enhancers^38^. However, how regulatory grammar emerges, when a trajectory commits to a productive regulatory program, and why trajectories become trapped in some cellular contexts remain unclear.

Here, we present <u>G</u>uided <u>O</u>ptimization of *<u>C</u>is*-<u>R</u>egulatory <u>E</u>lements (GO-CRE), a reinforcement learning framework for interpretable *in silico* generation of synthetic CREs. GO-CRE is built on HybriDNA^39^, an efficient decoder-only DNA language model with a hybrid Transformer–Mamba2 architecture that supports iterative sequence optimization under practical computational constraints. Rather than treating sequence generation as a black-box process, GO-CRE reconstructs the sequence-grammar landscape across reinforcement learning iterations using deconvolved k-mer and motif features. These features are then used to interpret trajectory dynamics and, when needed, guide biologically informed reward shaping.

Across K562, HepG2, and SK-N-SH cells, CRE optimization follows staged and directional trajectories comprising search, commitment, and optimization phases. Productive K562 and HepG2 trajectories cross local grammar barriers and converge on lineage-aligned regulatory programs, whereas SK-N-SH trajectories remain confined to a local basin and fail to produce robust target-selective activity. In HepG2, GO-CRE further identifies a low-complexity polyG-associated trap. Penalizing this feature redirects optimization toward hepatocyte-associated HNF/FOXA-centered motif programs, demonstrating that trajectory analysis can diagnose and correct unproductive optimization paths. Furthermore, generated CREs retain sequence diversity while converging on compact, lineage-aligned regulatory grammars. In HepG2, a recurrent HNF1B–FOXA1–HNF4A motif organization emerges progressively during RL and differs from the more heterogeneous architectures of endogenous STARR-seq-supported CREs. K562-generated CREs similarly enrich hematopoietic GATA/RUNX-associated programs. Fluorescence-based lentiviral massively parallel reporter assays validate target-selective activity in K562 and HepG2. HepG2-generated CREs show higher average reporter activity than endogenous CREs, while the strongest generated and endogenous elements achieve comparable activity through distinct motif architectures.

Collectively, GO-CRE shifts synthetic CRE design from endpoint-centered optimization toward a trajectory-resolved and steerable process. By linking iterative sequence changes to regulatory grammar and feeding interpretable features back into reward design, GO-CRE provides a framework for determining when optimization succeeds, why it fails, and how it can be redirected. These capabilities support both programmable regulatory sequence engineering and the systematic study of how sequence grammar gives rise to context-specific regulatory function, enabling GO-CRE to achieve future context-specific gene control applications.

## Results

### HybriDNA provides an efficient generative backbone for interpretable and tunable reinforcement learning

To dissect and optimize synthetic CRE generation using reward-based feedback, we introduced an interpretable and tunable reinforcement learning framework for context-specific regulatory sequence design (Figure 1A). Since iterative optimization requires repeated sequence sampling and policy updates, the framework requires a generative model that can operate under practical GPU memory constraints. We thus used HybriDNA, a decoder-only genomic language model that interleaves Mamba2 selective state-space blocks with Transformer attention layers at a 7:1 ratio, leveraging the sub-quadratic scaling of Mamba2 to process long genomic contexts efficiently while preserving nucleotide resolution via periodic attention layers. HybriDNA was trained and evaluated using a multispecies corpus comprising approximately 173 billion nucleotides from 851 species, including 816 training species and 35 held-out validation species, and in this study was subsequently further adapted with reverse-complement echo embeddings for sequence-understanding tasks (Figure 1B; Figure S1A-S1B).

**Figure 1.**
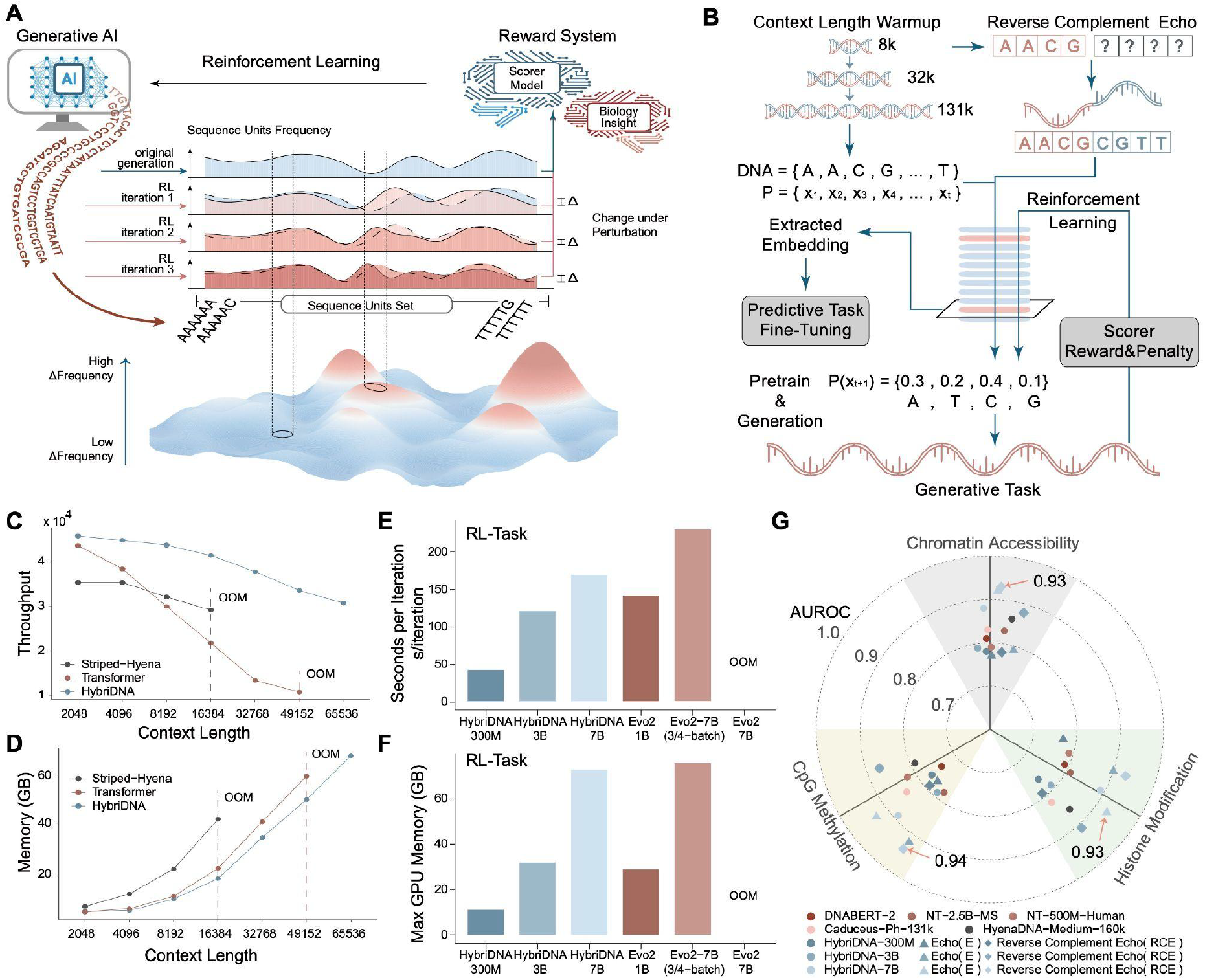
HybriDNA provides an efficient generative backbone for reinforcement learning-based CRE optimization. (A) Schematic of the GO-CRE concept. A generative DNA language model proposes candidate CREs, a predictor scores cell-type-specific activity, and reward feedback together with biological priors guides iterative sequence optimization and trajectory reconstruction. (B) Schematic of the optimized HybriDNA workflow. Three context lengths of DNA fragments, 8k, 32k and 131k, are input into the Transformer-Mamba2 hybrid architecture for pretraining at single nucleotide resolution, and then fine-tuned with reverse-complement echo embedding (RCE) approach. Mamba2 selective state-space blocks and Transformer decoder layers are interleaved at a 7:1 ratio. For generation tasks, reinforcement learning is employed to enhance the generative DNA specifically. (C–D) Throughput (Tokens per second) (C) and GPU memory usage (D) of HybriDNA, pure Transformer, and StripedHyena across input context lengths using models with matched parameter counts (300M). x-axis labeled input DNA context length. OOM, out of memory. (E–F) Iteration time (E) and maximum GPU memory usage (F) for RL tasks comparing HybriDNA and Evo2 models on a single 80G A100 GPU. HybriDNA tested on 300M, 3B and 7B versions, Evo2 tested on 1B and 3/4 batched 7B versions. Full-batch Evo2-7B exceeded the 80-GB memory limit and is shown as OOM. (G) Performance on the BEND benchmark shown as AUROC for chromatin accessibility, histone modification, and CpG methylation prediction across genomic language models. For HybriDNA, circles denote the initial model, triangles denote the echo model, and squares denote the reverse-complement echo model.

At a matched scale of approximately 300 million parameters and under identical single-GPU settings, HybriDNA achieved higher training throughput and lower peak memory consumption than a pure Transformer and StripedHyena across all tested context lengths, and the advantage increased with sequence length. (Figures 1C-1D). We further tested efficiency during reinforcement learning. Under matched PPO settings on a single 80-GB A100 GPU, HybriDNA-7B completed full-batch optimization, whereas Evo2-7B exceeded the memory limit. HybriDNA-7B required an iteration time comparable to Evo2-1B, and the 300M and 3B HybriDNA models were faster than Evo2-1B (Figures 1E-1F).

We next evaluated the genomic sequence understanding capacity of HybriDNA. On the BEND benchmark, HybriDNA-7B achieved strong performance across chromatin accessibility, histone modification and CpG methylation prediction tasks, outperforming or matching established genomic language models including DNABERT2, Nucleotide Transformer, HyenaDNA and Caduceus variants (Fig. 1G). HybriDNA also performed strongly across the Genome Understanding Evaluation tasks, including transcription-factor binding, promoter and core-promoter detection, epigenetic-mark prediction, viral-variant classification, and splice-site recognition (Supplementary Figure 1C-1I).

### GO-CRE reconstructs and redirects sequence-grammar trajectories during CRE optimization

To extend the efficient RL capacity of HybriDNA toward cell-type-specific CRE design, we developed <u>G</u>uided <u>O</u>ptimization of *<u>C</u>is*-<u>R</u>egulatory <u>E</u>lements (GO-CRE), a closed-loop framework that reconstructs sequence-grammar trajectories during RL and uses them to steer sequence optimization. This design makes CRE generation interpretable and tunable by feeding deconvolved grammar features back into biologically informed reward shaping (Figure 2A). GO-CRE uses an MPRA-finetuned HybriDNA model as the sequence generator and a pretrained regulatory activity predictor from CODA^36^ to score each candidate sequence. For each generated sequence, the reward is defined as the activity in the target cell type minus the maximum activity across off-target cell types (MinGap score), thereby favoring high regulatory activity in the target cell type while simultaneously penalizing activity in non-target cell types (Figure 2A, left).

**Figure 2.**
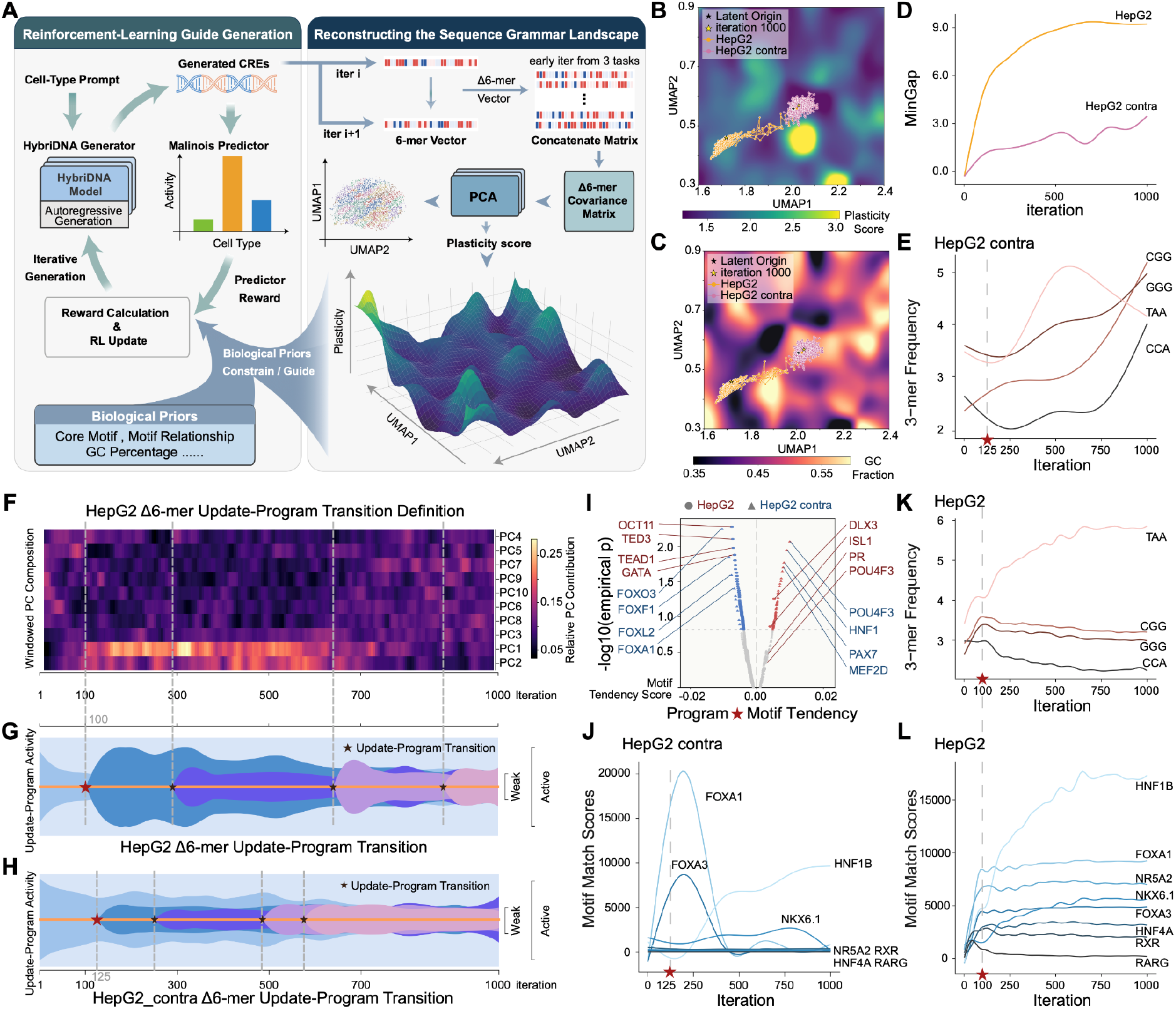
GO-CRE reconstructs interpretable and tunable sequence-grammar trajectories. (A) Detailed overview of the GO-CRE framework. Left: Closed-loop reinforcement learning (RL) workflow for cell-type-specific CRE generation using predictor-based rewards and biological priors. Right: Sequence-grammar landscape reconstruction from Δ6-mer changes across RL iterations, followed by feedback of deconvolved grammar features to guide subsequent rounds of optimization. (B–C) Optimization trajectories of polyG-penalized HepG2 sequences (HepG2, orange) and unpenalized controls (HepG2 contra, pink) projected onto the reconstructed grammar landscape colored by plasticity score (B) or GC fraction (C).Iteration 1000 labeled as star in the map. (D) MinGap score (see Methods) trajectories across RL iterations for HepG2 (orange) and HepG2 contra (pink). (E) Representative 3-mer frequency trajectories during unpenalized HepG2 optimization (HepG2 contra). Gray dashed lines mark iteration 125. (F) Windowed principal component composition used to define Δ6-mer update programs across HepG2 iterations. Relative principal component contributions are shown at right. (G-H) Activities of inferred update programs across HepG2(G) and HepG2 contra(H) RL iterations. Stars indicate the emergence of new update programs. Red star marks iteration 100(G) or 125(H). (I) Corrected motif tendency scores for the first newly emerged update program in HepG2 (red) and HepG2 contra (blue). (J) Representative motif match score trajectories during unpenalized HepG2 optimization (HepG2 contra). Gray dashed lines mark iteration 125. (K–L) Representative 3-mer frequency trajectories (K) and motif match score trajectories (L) during HepG2 optimization. Gray dashed lines mark iteration 100, corresponding to the first major update program. For panels D–H, and J–L, the x axis indicates RL iteration (1–1,000), increasing from left to right; y axes indicate the corresponding plotted features.

To make GO-CRE interpretable and tunable, we deconvolved each RL iteration into k-mer and motif-level features. We consider that the fine-tuned HybriDNA generator defines an underlying sequence grammar landscape that constrains accessible regulatory solutions, whereas RL progressively redistributes generated sequences across this landscape during optimization. Earlier iterations are therefore expected to remain more reflective of inherited grammar structure and provide a suitable window for recovering its dominant organization. Under this framework, sequence sets are represented by k-mer frequency profiles, and Δk-mer frequencies between consecutive iterations are used as proxies for local displacements on the grammar landscape. Guided by non-negative matrix factorization (NMF) of motif trajectories, we integrate the first 40 Δk-mer profiles from three tasks to construct a covariance matrix (Figure S2A–S2C). Considering the tradeoff between feature dimensionality and empirical effective rank, we selected 6-mers as the fundamental units and used principal component analysis (PCA) on the resulting covariance matrix to resolve the dominant axes of sequence grammar variation and reconstruct the grammar landscape (Figure S2D–S2E). The reconstructed grammar space further revealed biologically meaningful structure. For example, GC content was correlated with differential sequence plasticity under RL perturbation, with low-GC features exhibiting greater plasticity than GC-rich features (Figure S2E–S2G). Such deconvoluted information was then incorporated into reward calculation to tune the RL process and mitigate potential generative traps (Figure 2B).

We next tested whether trajectory-derived grammar features could be used to actively tune the generation process. We take HepG2 as a representative cell line, and project the original generative RL process into the sequence-grammar landscape. The unpenalized HepG2 trajectory (HepG2 contra) became trapped in a high-GC local basin of the grammar landscape (Figure 2B-2C), and the MinGap score continued to increase across iterations without rapid saturation (Figure 2D). With further 3-mer feature dissection, we found that this trap-prone trajectory was accompanied by continuous accumulation of polyG features in generated sequences (Figure 2E). Thus, we introduced a polyG penalty to the reward function, and redirected reinforcement learning away from the low-complexity regime(Figure 2B-2C). This intervention produced an earlier saturation and more sustained improvement in MinGap score across the RL iterative generation process (Figure 2D), demonstrating that a trajectory-derived feature could be fed back into reward design.

With RL iterations mapping onto the sequence-grammar landscape, we defined three phases of the RL generation process, which we termed search, commitment, and optimization (Figure 2B–2C). During the search phase, generated sequences broadly explore grammar space while sampling features compatible with the reward function. As reward-aligned features accumulate, trajectories enter a commitment phase marked by directional movement across grammar barriers and enrichment of target-associated features. When the reward approaches saturation, the RL process stabilizes in the optimization phase to gain diverse features without changing the core grammar space position. These three phases of generation are further captured by the program transitions over iterations. In polyG-penalized HepG2 optimization, we identified the major program shifts in 1) iteration 100, mainly referring to “search->commitment” transition, 2) iteration 290, representing an intermediate state in which a new update program emerged while the commitment-associated program remained predominant and 3) iteration 640, mainly referring to accomplishment of “commitment->optimization” transition with the emergence of a new dominant program(Figure 2F). Compared with the unpenalized HepG2 contra trajectory, the polyG-penalized trajectory maintained stable phase transitions in which newly emerged programs gained sustained dominance, particularly in “search->commitment” transition point (Figure 2G-2H), further demonstrating the importance of efficient sequence feature commitment during CRE generative tuning.

We further dissected the biological logic underlying RL optimization by moving from sequence-level programs to motif-level interpretation. Sparse program weights in PC space were mapped back to signed 6-mer preferences and then integrated over motif 6-mer profiles to infer program-associated motif biases on the search->commitment transition point (Figure 2I, S2H, see Methods). In the unpenalized HepG2 contra group, two major lineage-specific transcription factor family motifs, FOXA and HNF, show temporally offset waves of enrichment along the optimization trajectory(Figure 2J, S2I). With polyG penalization, HNF1B emerged as a dominant motif without loss of FOXA-family scores, and the second wave of HNF1B enrichment coincided with the iteration 100 transition point(Figure 2K-2L, S2J). These results indicate that polyG penalization promoted stable co-occurrence of FOXA- and HNF-associated motif programs in generated sequences.

Together, these analyses show that GO-CRE converts reinforcement learning from an endpoint-only optimization into an observable sequence-grammar trajectory and uses that trajectory to diagnose and correct an unproductive solution.

### GO-CRE generates cell-type-specific CREs with distinct regulatory programs

We next asked whether GO-CRE could generate cell-type-specific CREs across multiple cellular contexts. We projected the generation trajectories for HepG2, K562, and SK-N-SH onto the reconstructed sequence-grammar landscape and quantified step size and directional straightness across iterations to characterize search, commitment, and optimization phases across cell lines (Figure 3A-3B, S3A-S3B). Similar to HepG2, both K562 and SK-N-SH acquired transcription factor motif features during optimization, accompanied by phase-associated transition points (Figure 3C-3D, S3C-S3D). Particularly, K562 optimization progressively acquired hematopoietic regulators, including GATA1- and RUNX-associated motifs (Figure 3C), and its Δ6-mer trajectory contained multiple stable update-program transitions (Figure S3C), complementing the HNF/FOXA-centered program identified in HepG2 and supporting the progressive acquisition of lineage-associated features during RL process. By contrast, SK-N-SH showed weaker convergence on neural-lineage grammar and stronger enrichment of NF-κB and other inflammatory or broadly active transcription-factor programs (Figure 3D, S3D), which is associated with SK-N-SH remaining confined to a high-GC basin after the search phase (Figure 3A-3B). As MinGap scores increased or stabilized across the three cell types, sequence diversity, measured by normalized 6-mer Vendi score, also approached a stable level during the optimization phase (Figure 3E-3F). While HepG2 and K562 generated CREs stabilized at diversity levels comparable to Malinois-filtered natural sequences, SK-N-SH retained some sequence diversity but did not follow a productive three-phase trajectory toward a lineage-aligned regulatory program (Figure 3A, 3F), suggesting an efficient generation tuning process can actually maintain, rather than reduce, sequence diversity of the synthetic CREs.

**Figure 3.**
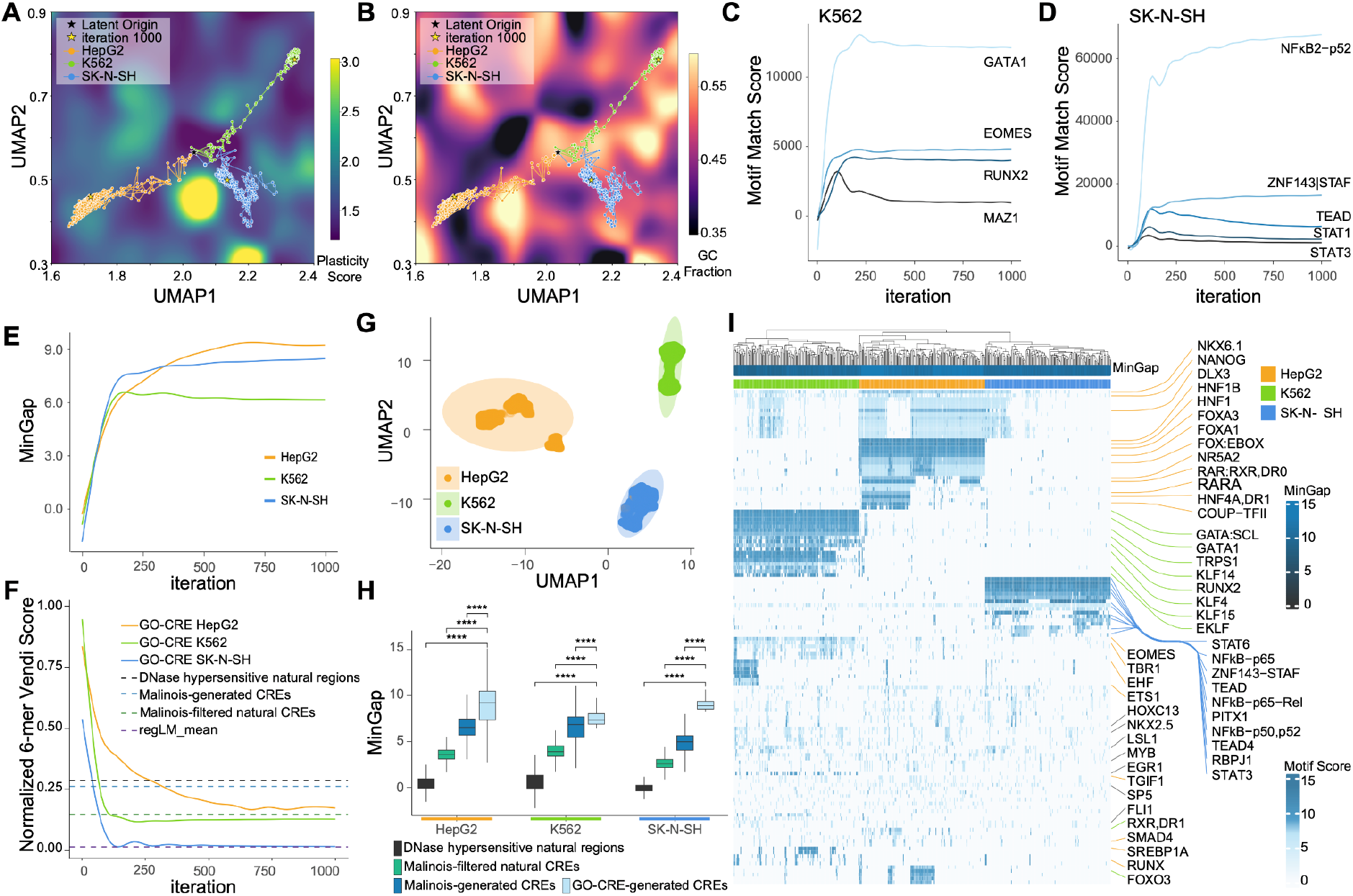
Cell-type-specific optimization converges on distinct regulatory programs. (A–B) RL trajectories projected onto the reconstructed sequence-grammar landscape and colored by plasticity score (A) or GC fraction (B). The latent origin corresponds to the centroid of iteration 1 across tasks. Trajectories over 1,000 RL iterations are shown for HepG2 (yellow), K562 (green), and SK-N-SH (blue). Iteration 1000 labeled as star in the map. (C–D) Representative motif match score trajectories during K562 (C) and SK-N-SH (D) optimization. (E) MinGap trajectories across RL iterations for HepG2 (yellow), K562 (green), and SK-N-SH (blue). (F) Normalized 6-mer Vendi score across 1,000 RL iterations for GO-CRE HepG2 (yellow), GO-CRE K562 (green), and GO-CRE SK-N-SH (blue). Dashed lines show the normalized 6-mer Vendi scores averaged across three cell types for sequence sets from DNase hypersensitive natural regions (black), Malinois-generated CREs (blue), Malinois-filtered natural CREs (green), and regLM-generated CREs (purple). (G) UMAP visualization of 9,000 final generated CREs from each target cell line: HepG2, K562, and SK-N-SH. (H) MinGap scores of GO-CRE-generated CREs, DNase hypersensitive natural regions, Malinois-filtered natural CREs, and Malinois-generated CREs across HepG2, K562, and SK-N-SH. Statistical significance was assessed using two-sided Wilcoxon rank-sum tests. GO-CRE was compared to each control group within each cell type. **** *p* < 0.0001. (I) Heatmap of motif scores across representative transcription factor motifs in final generated CREs from HepG2, K562, and SK-N-SH. Columns represent individual generated CREs (n = 150 per cell type), ordered by hierarchical clustering. Post-evolved CREs were selected from the top 5 MinGap clusters, with the top 30 sequences from each cluster. Rows correspond to distinct motifs (n = 132). Top annotation bars indicate MinGap and target cell type.

The endpoint generated synthetic CREs segregated clearly across the three target cell lines (Figure 3G), and exhibited increased predicted regulatory activity compared to CREs generated by the CODA and regLM models across all three cell lines, as well as K562- and HepG2-specific CREs generated by DNA-Diffusion (Figure 3H and Figure S3E–S3F). The generated CREs retained substantial diversity while displaying pronounced cell-line-specific transcription factor (TF) motif features within each lineage (Figure 3I, Figure S3G–S3L). In each cell line, we found some motifs were nearly universal within a lineage, such as FOXA1 and HNF1B in HepG2, and GATA1 in K562. Other motifs, such as RARA in HepG2, were enriched only in specific subclusters (Figure 3I). These results indicate that GO-CRE captures core lineage requirements while avoiding collapse onto a single repeated sequence family.

Together, these results show that GO-CRE generates synthetic CREs that are diverse and enriched in biologically meaningful features. Successful outputs in HepG2 and K562 combine strong predicted activity with lineage-aligned motif programs, whereas SK-N-SH provides a contrasting case in which diversity is retained but productive lineage convergence is weaker.

### GO-CRE-generated and endogenous CREs encode distinct motif organizations

We next compared the motif architecture of GO-CRE-generated CREs with that of endogenous CREs. Whereas endogenous CREs showed broad motif heterogeneity, GO-CRE-generated CREs formed one or two dominant motif clusters in HepG2 and K562 (Figure S4A–S4D), suggesting convergence on shared higher-order regulatory grammars despite preserved sequence diversity. Orthogonal ChromBPNet models predicted that cell-type-specific designs were more accessible in matched than unmatched cell types (Figures 4A-4B). Direct comparison with endogenous STARR-seq-supported CREs further showed stronger lineage-aligned motif enrichment in the generated CREs. In HepG2, HNF1B, HNF4A, FOXA1, and other hepatocyte-associated motifs were preferentially enriched, whereas K562-generated CREs concentrated motifs of hematopoietic lineage factors including GATA1/2, RUNX2, and EOMES (Figure 4C–4D). These results indicate that GO-CRE distilled compact lineage-associated motif programs rather than recapitulating the full heterogeneity of endogenous CREs.

**Figure 4.**
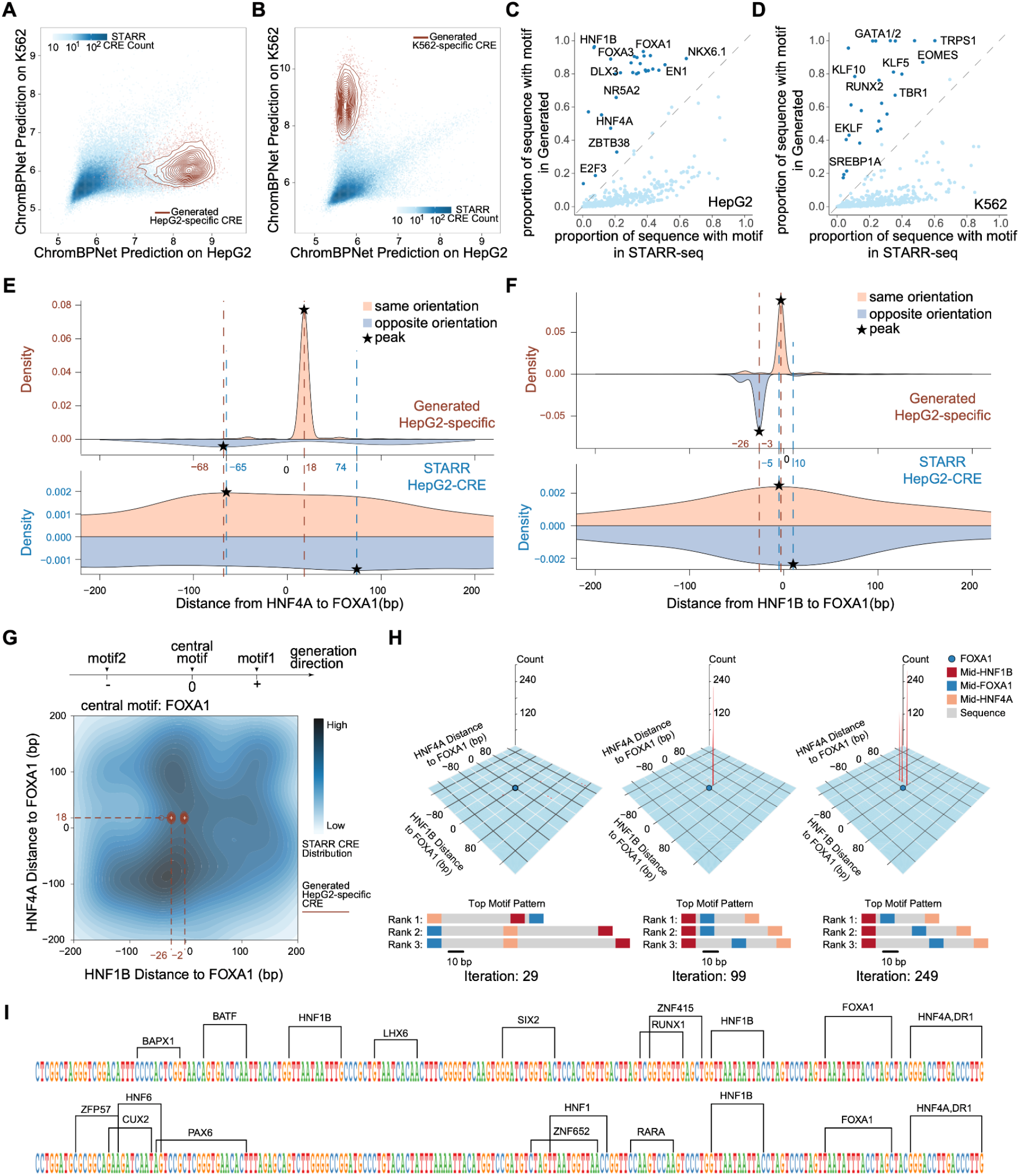
Generated and endogenous CREs encode distinct motif organizations. (A–B) Density projections of generated HepG2-specific (A) or K562-specific (B) CREs in ChromBPNet prediction space defined by predicted accessibility in HepG2 and K562. Endogenous STARR-seq CRE distributions are shown as background density. (C–D) Comparison of motif prevalence in generated and endogenous STARR-seq CREs from HepG2 (C) and K562 (D). Each point represents one motif, plotted as the proportion of sequences containing that motif in generated CREs versus endogenous CREs. (E–F) Density plots of motif spacing centered on FOXA1 in generated HepG2-specific and endogenous STARR-seq HepG2 CREs. Distances from HNF4A to FOXA1 (E) and from HNF1B to FOXA1 (F) are shown. Positive and negative densities indicate same and opposite motif orientations, respectively. Asterisks mark peak positions. (G) Two-dimensional density of HNF1B-to-FOXA1 and HNF4A-to-FOXA1 distances centered on FOXA1 in endogenous STARR-seq HepG2 CREs (blue) and generated HepG2-specific CREs (red). (H) Iterative emergence of HNF1B–FOXA1–HNF4A motif organization during HepG2 optimization. Distance distributions relative to the central FOXA1 motif and top-ranked motif patterns are shown for representative iterations 29, 99, and 249. Full process see supplementary video 1. (I) Motif organization of two representative generated HepG2-specific CREs.

We next tested whether this convergence extended beyond motif abundance to motif syntax. In HepG2, distances from HNF4A and HNF1B to a central FOXA1 motif showed sharper, recurrent peaks in generated CREs than in endogenous CREs, with orientation-specific structure evident on both sides of FOXA1 (Figures 4E-4F). Joint analysis of the two distances revealed a concentrated HNF1B-FOXA1-HNF4A configuration in generated sequences, compared with the more diffuse organization of endogenous CREs (Figure 4G).

This higher-order organization emerged progressively during reinforcement learning. Across representative iterations 29, 99, and 249, the HNF1B-FOXA1-HNF4A distance distribution became increasingly structured, and the full trajectory showed continued consolidation of the motif phrase (Figure 4H; Supplementary Video 1). Representative endpoint designs contained compact arrangements of the three motifs with recurrent spacing and orientation (Figure 4I). Thus, GO-CRE-generated sequences exhibited cell-type-biased regulatory predictions while using more condensed motif organization than matched endogenous CREs.

### LentiMPRA validates context-specific activity of generated synthetic CREs

We next used a fluorescence-based lentiviral massively parallel reporter assay (MPRA) to validate the activity and specificity of GO-CRE-generated enhancers. We designed a dual-reporter vector in which hPGK-driven mCherry served as an internal normalization control for viral copy number and integration effects, whereas candidate CREs were cloned upstream of a minimal promoter-GFP cassette to report enhancer activity. mCherry-positive cells were then sorted into four bins according to the GFP/mCherry ratio, representing increasing levels of enhancer activity (Figure 5A, Figure S5A–S5B).

**Figure 5.**
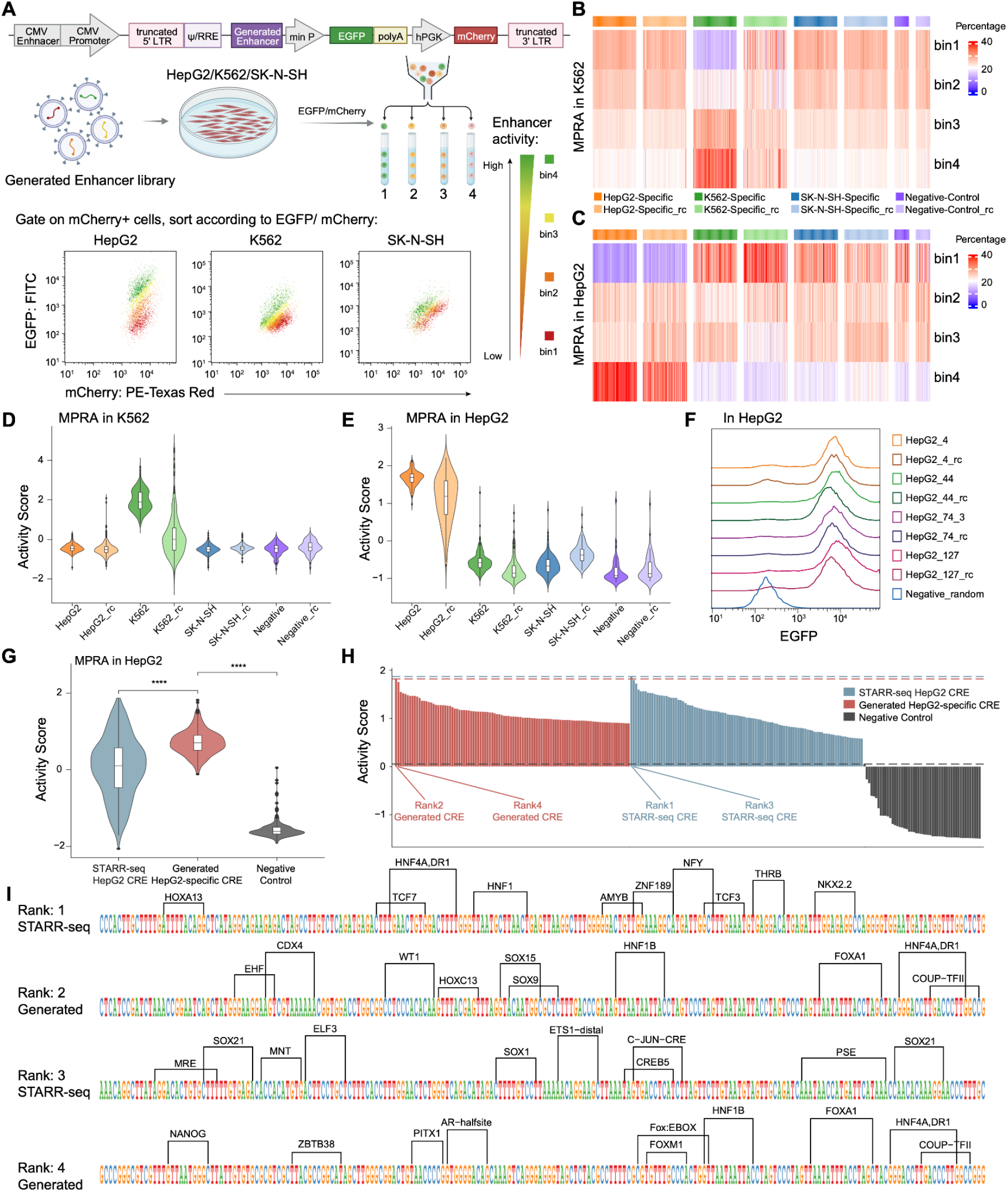
LentiMPRA validates cell-type-specific activity of generated CREs. (A) Schematic of the fluorescence-based LentiMPRA workflow. Candidate CRE libraries were cloned upstream of a minimal promoter-EGFP reporter in a dual-color lentiviral vector containing hPGK-mCherry as an internal normalization control. Lentiviral libraries were transduced into HepG2, K562, or SK-N-SH cells. At 4 days post-transduction, mCherry-positive cells were sorted into four bins according to the EGFP/mCherry ratio (see Figure S5), and CRE abundance in each bin was quantified by amplicon sequencing. (B–C) Heatmaps of CRE enrichment across bins 1–4 in K562 (B) and HepG2 (C) cells. Columns represent individual CREs, and rows represent sorted bins of increasing EGFP/mCherry ratio as defined in (A). Top annotation bars indicate CRE class. (D–E) LentiMPRA activity scores for all tested CREs in K562 (D) and HepG2 (E) cells, shown as violin plots. (F) Representative EGFP fluorescence histograms from individual validation of HepG2-generated CREs. (G) LentiMPRA activity scores in HepG2 for endogenous CREs identified by STARR-seq and generated HepG2-specific CREs selected by ChromBPNet, shown as violin plots. \*\*\*\**p* < 0.0001. (H) Ranked activity scores of the top 25% most active endogenous and generated HepG2 CREs measured by LentiMPRA. Representative sequences shown in (I) are indicated. (I) TF motif organization of the four highest-ranking endogenous and generated HepG2 CREs shown in (H). For panels B–H, RC denotes reverse-complemented sequence. Negative-control sequences were derived from GM12878-accessible regions lacking accessibility in the indicated target cell type. Data represent 2–3 independent experiments.

We next profiled generated CRE libraries from HepG2, K562, and SK-N-SH together with reverse-complement and negative controls. In K562, K562-specific generated CREs were preferentially enriched in the high-activity bins and showed higher activity scores than off-target generated sequences or reverse-complement controls (Figure 5B and 5D). Similarly, in HepG2, HepG2-specific generated CREs were concentrated in the most active bins and outperformed off-target generated sequences and controls (Figure 5C and 5E). By contrast, SK-N-SH-generated CREs did not show clear target-selective activity, consistent with the trap-prone optimization behavior observed in the sequence-grammar landscape (Figure S5C). Clonal validation of individual HepG2-generated CREs further confirmed strong HepG2-specific reporter activation relative to a random-sequence negative control (Figure 5F).

Finally, we compared post-evolved HepG2-generated CREs with endogenous HepG2 STARR-seq CREs. On average, generated CREs showed higher MPRA activity than endogenous CREs, although the top-performing sequences from the two groups reached comparable activity levels (Figure 5G–5H). Despite comparable activity among the strongest sequences, the underlying motif grammar remained distinct: the highest-ranking generated CREs preserved an HNF1B–FOXA1–HNF4A-like organization, whereas top endogenous CREs displayed more heterogeneous motif combinations (Figure 5I). These data indicate that GO-CRE can generate diverse, cell-type-specific synthetic enhancers that are functionally competitive with endogenous regulatory elements while converging on concentrated lineage-aligned grammar features.

Taken together, our results establish GO-CRE as an interpretable and tunable framework for RL-guided synthetic CRE generation. By coupling efficient DNA language-model-based generation with grammar-level trajectory reconstruction and biologically guided reward shaping, GO-CRE enables the design of diverse enhancers with experimentally validated cell-type specificity. The contrast between successful optimization in K562 and HepG2 and failed optimization in SK-N-SH further indicates that synthetic CRE designability depends on whether RL trajectories can escape local grammar traps and converge on productive regulatory programs. These findings provide a foundation for trajectory-aware cis-regulatory design and for understanding how regulatory grammar emerges during synthetic sequence optimization.

## Discussion

Synthetic *cis*-regulatory elements offer a means to program context-specific gene expression for both basic research and gene and cell therapy applications^13,40,41^. Generative models can expand the accessible design space beyond endogenous regulatory repertoires, particularly for cellular states that are poorly represented in available datasets. However, most sequence design approaches remain focused on endpoint performance. Although predictive models can rank candidate sequences, they provide limited insight into how regulatory features emerge during optimization, why particular trajectories succeed or fail, or how biological knowledge can be used to correct an unproductive search. Consequently, score-driven optimization may exploit low-complexity or non-physiological sequence features that satisfy the predictor without establishing a coherent regulatory program.

GO-CRE addresses this limitation by treating synthetic CRE design as an observable and steerable process. Enabled by the computational efficiency of HybriDNA, GO-CRE supports iterative sequence generation while reconstructing changes in sequence grammar throughout optimization. The resulting trajectories resolve into search, commitment, and optimization phases, during which broad exploration is progressively narrowed toward stable regulatory programs. The combination of iterative sequence variation, functional selection, staged feature acquisition, and reward-dependent trajectory redirection supports viewing this process as an in silico analogue of directed evolution. In this framing, the emphasis is not only on selecting a functional endpoint, but also on understanding and controlling the path through which that endpoint is reached.

The resulting trajectories reveal that successful synthetic CRE design depends on the path through sequence-grammar space. Productive optimization was associated with coordinated transitions between sequence-update programs and convergence on lineage-aligned regulatory features. The polyG intervention in HepG2 provides a proof of principle that such failure modes can be identified during optimization and converted into optimization constraints. Thus, GO-CRE functions as both a design engine and a diagnostic framework for CRE generation.

The generated CREs illustrate how synthetic and endogenous regulatory elements may reach similar functions through different grammatical solutions. GO-CRE designs retained sequence diversity but concentrated lineage-associated motifs into comparatively compact regulatory programs. In HepG2, the progressive emergence of an HNF1B-FOXA1-HNF4A organization suggests that higher-order motif syntax can arise during iterative optimization rather than appearing only as an endpoint enrichment. The earlier emergence of FOXA-family motifs relative to HNF1B is compatible with established roles of FOXA factors as pioneer regulators and HNF factors in hepatocyte identity^42, 43^. This correspondence is intriguing, although it should not be interpreted as evidence that the optimization trajectory directly recapitulates liver development.

Lentiviral MPRA confirmed target-selective activity for K562 and HepG2 designs and showed that highly active synthetic and endogenous CREs can achieve comparable reporter output despite distinct motif organizations. The weaker SK-N-SH result further indicates that trajectory analysis may help identify cellular contexts in which the available training data, predictor or reward formulation is insufficient for productive design. However, reporter assays measure autonomous enhancer activity outside native genomic and chromatin environments. Future work should therefore test GO-CRE designs at endogenous loci, in primary cells and in vivo, and should incorporate state-resolved chromatin and transcriptional information into the optimization objective.

Several future directions emerge from this work. Extending GO-CRE beyond reporter-based optimization will be important for matching synthetic CRE design to endogenous chromatin and genomic context, particularly because reporter-active and native regulatory grammar may not fully overlap. Broadening the optimization objective to incorporate chromatin-aware and stimulus-responsive signals should further expand the design space. In addition, the weaker performance in SK-N-SH highlights the need for richer cell-state-resolved training data and priors^44,45^, especially for heterogeneous cellular systems. Future versions of GO-CRE should therefore integrate matched functional and chromatin datasets, including state-resolved accessibility and transcriptional information, together with more flexible search strategies that preserve alternative trajectories rather than converging prematurely on a single dominant solution.

Overall, these findings establish GO-CRE as a framework for making CRE design interpretable and steerable. By exposing the trajectories through which regulatory grammar is acquired, GO-CRE allows unproductive solutions to be recognized and corrected during optimization rather than only after sequence generation is complete. This shift from black-box sequence optimization to trajectory-aware regulatory design provides a foundation for controllable CRE engineering and for systematic analysis of regulatory grammar dynamics.

## Resource availability

### Lead Contact

Further information and requests for resources and data should be directed to and will be fulfilled by the lead contact, Dr. Zeyu Chen.

### Data availability

Publicly available MPRA datasets used for HybriDNA fine-tuning were obtained from ENCODE, and accession numbers together with detailed information are provided in Supplementary Table 1. Additional public datasets used in this study are listed in the deposited data section of the Key Resources Table. MPRA datasets generated in this study are available on GSE330487.

### Code availability

The implementation of the HybriDNA model, the GO-CRE framework, scripts for sequence grammar landscape analysis, and code for MPRA experimental data processing are available at https://github.com/zeyuChenLab/GO-CRE.git.

## Acknowledgements

We thank Ting Han’s lab at NIBS for providing HepG2, K562 and 293T cell lines, Salvador Casaní-Galdón, Yanjing Liu, Shuang Li, Ruochi Zhang, and Bradley E. Bernstein for initial idea shaping. Jiecong Lin, Bohan Li, Han Yuan, Zhijian Li, Xi Dawn Chen and Fadi Najm for helpful discussions. This work is supported by the “U35 Pei Miao Funding Program” of Changping District, Beijing (No. CHPU35202501001), Beijing Municipal Science & Technology Commission (No.5264034, No.Z231100004823015), Noncommunicable Chronic Diseases-National Science and Technology Major Project (No. 2023ZD0501204),Open Funding for National Center for Protein Sciences at Peking University (No.KF-202501) and Zhongguancun Academy (No. XTS0010).

## Author Contributions

M.M., W.B., and Z.C. conceived the study. M.M. and G.L. supported the use of HybriDNA with input from P.D., Q.Y., C.C., H.L., and T.Q. M.M., W.B., and Z.C. optimized the RC-echo embedding strategy and designed the GO-CRE reinforcement learning framework and biological-prior discriminator. W.B. performed bioinformatic analyses and scoring with support from Z.C. and Y.H. Y.L., S.L., Z.Z., and S.Y. performed wet-lab experiments with support from Z.C. and M.X. P.D. and P.J. provided AI modeling and computational assistance with support from H.L. and T.Q. Z.C., W.B., M.M., Y.L., and S.L. wrote and edited the manuscript with assistance from Q.H., Y.Z., C.Z., M.X., Y.H., and T.Q.

## Declaration of generative AI and AI-assisted technologies in the manuscript preparation process

During the preparation of this work the authors used Codex in order to correct the coding, and GPT-5.4 / 5.5 Pro in order to correct the manuscript. After using this tool/service, the authors have reviewed and edited the content as needed and take full responsibility for the content of the published article.

## STAR Methods

### Key Resources Table

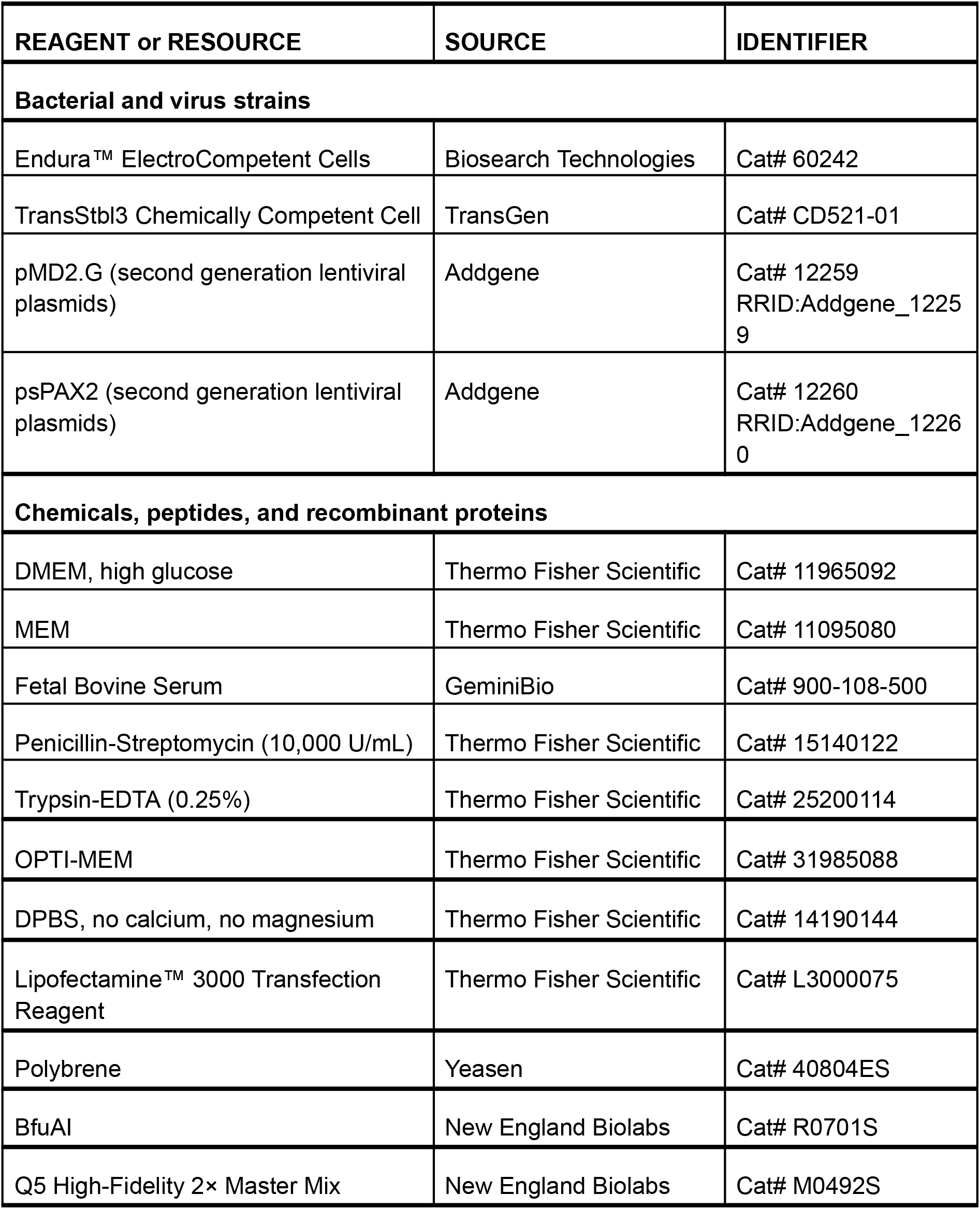
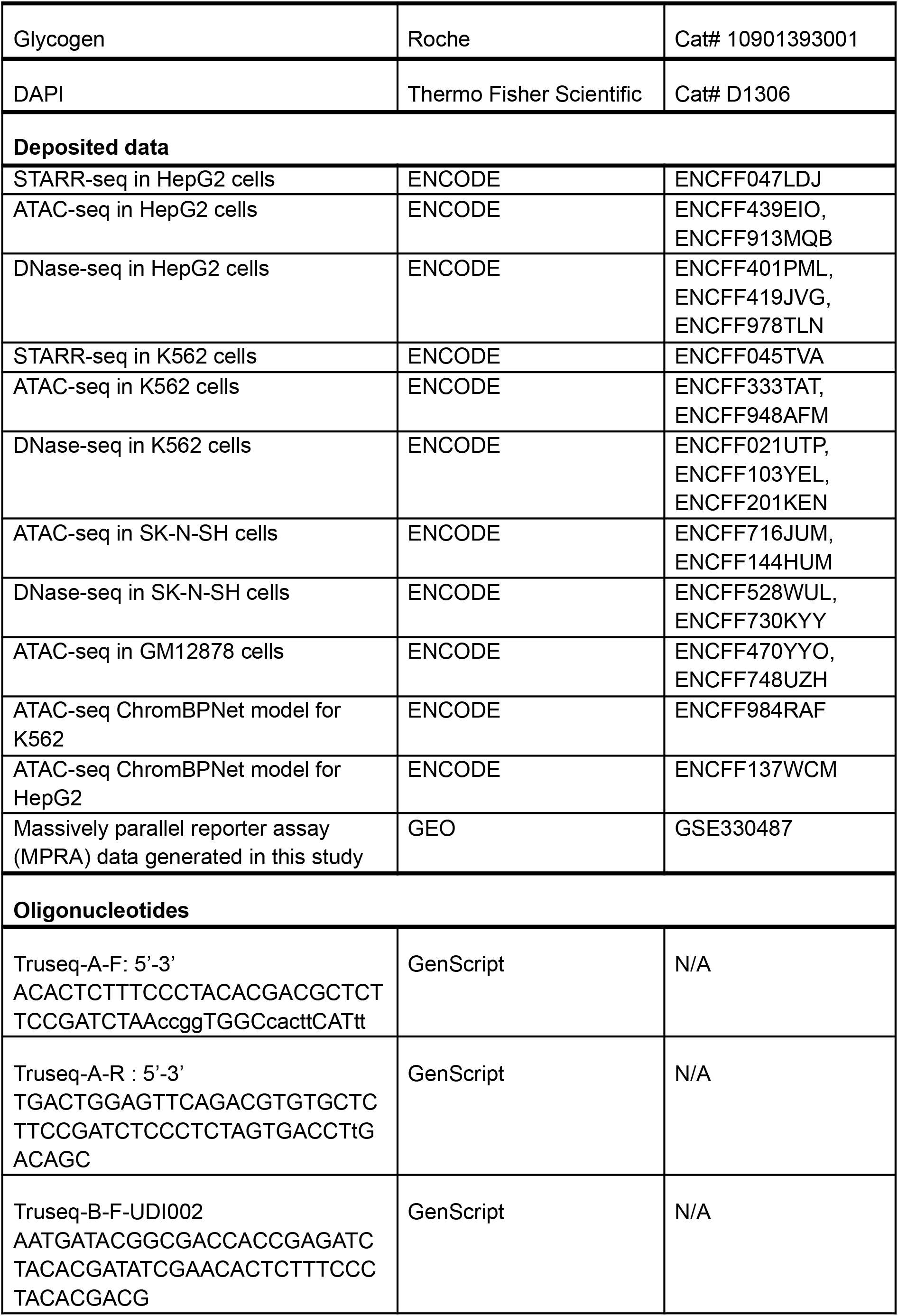
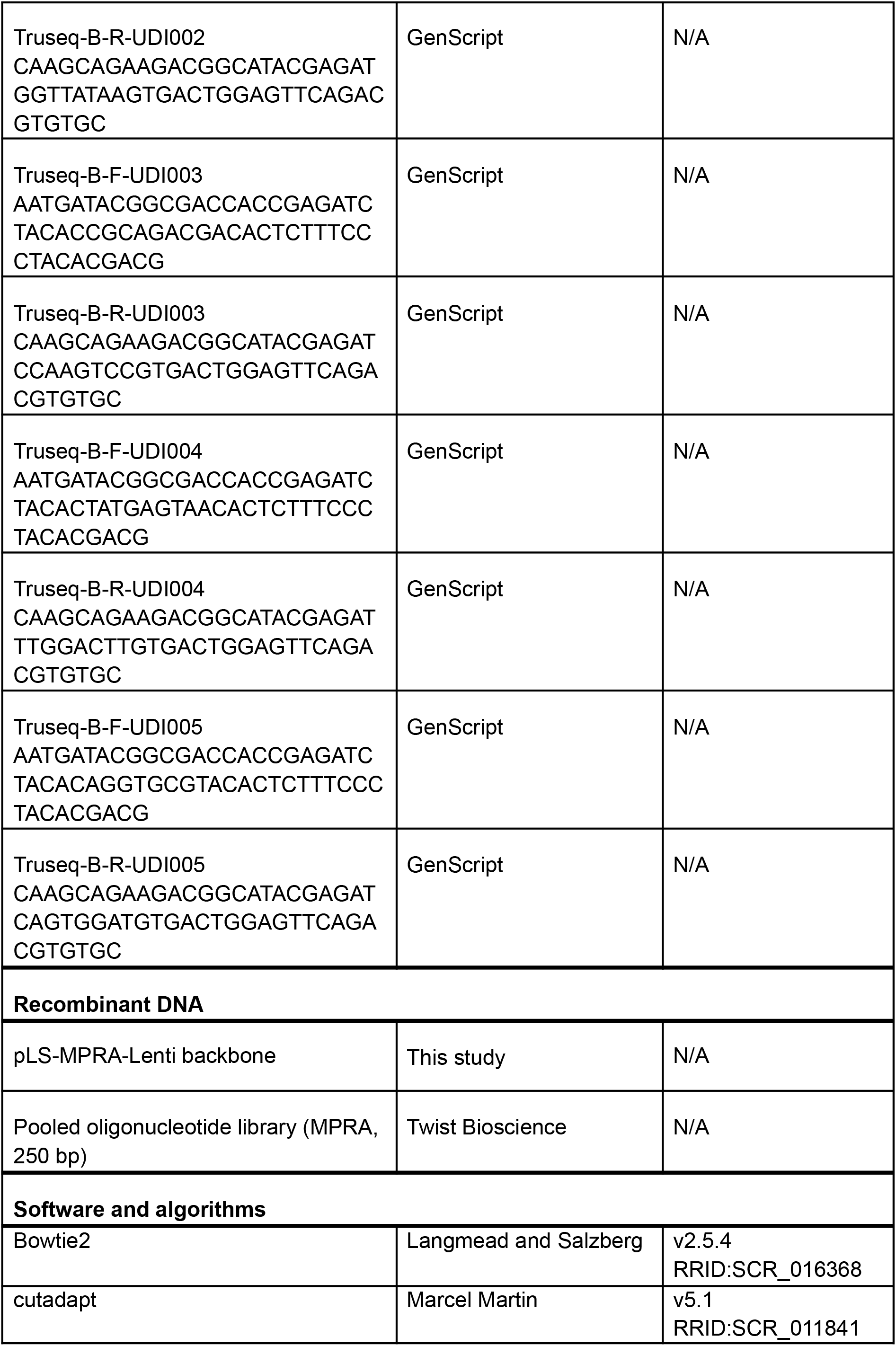
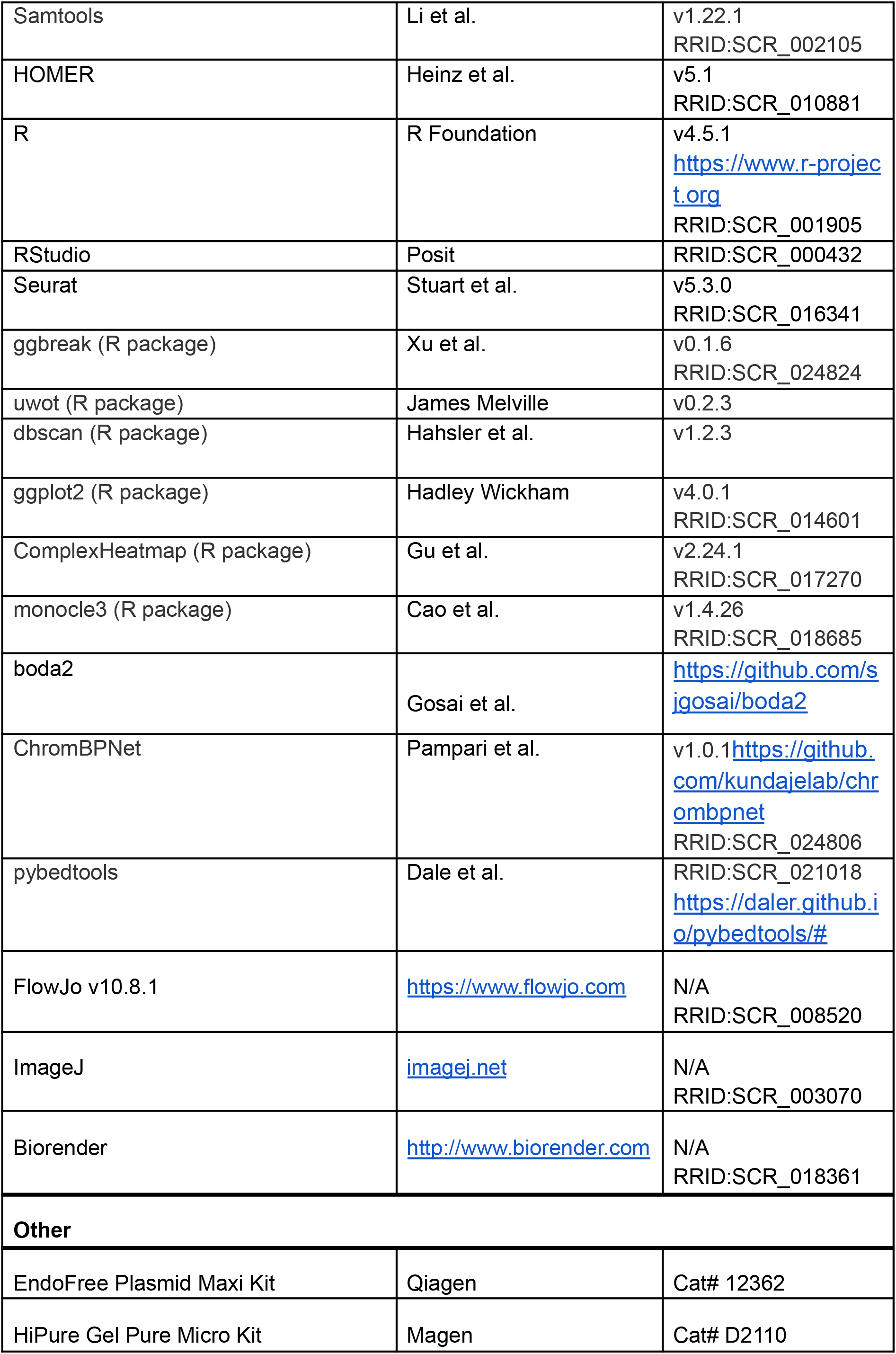
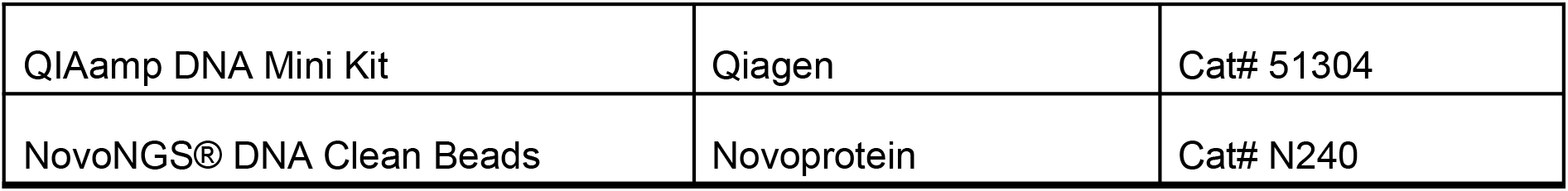

### HybriDNA Model Architecture

HybriDNA is a decoder-only autoregressive DNA language model that interleaves Mamba2 selective state-space model (SSM) blocks with Transformer attention blocks at a 7:1 ratio, designed to balance efficient long-range processing with single-nucleotide resolution. The Mamba2 blocks contribute sub-quadratic O(NL) complexity for long-range dependencies, where N is the SSM state dimension and L the sequence length, while the periodically interleaved Transformer blocks (causal masking, RMSNorm, group-query attention, feed-forward MLP) provide high-fidelity nucleotide-level interactions. Because the first layer is always a Mamba2 block, no explicit positional embeddings are required: positional information is implicitly encoded through the recurrent state dynamics of the SSM layers. Each Mamba2 block follows the State-Space Duality formulation^46^ with input-dependent parameter generation; full implementation details (block composition, normalization, gating) follow our previous description of HybriDNA^39^. The 7:1 hybrid ratio was empirically selected over a pure-Mamba2 baseline based on lower pretraining loss at matched parameter count, consistent with recent observations across hybrid language models^39,47–49^. We trained three variants of increasing scale to support downstream embedding and reinforcement learning workloads:

#### Model Variants

We trained three model variants to investigate scaling behavior:

| Variant | Parameters | Layers | Hidden Dim | Intermediate Dim | Attention Heads | KV Heads | Head Dim |
| --- | --- | --- | --- | --- | --- | --- | --- |
| HybriDNA-300M | 300M | 24 | 1024 | 2048 | 32 | 8 | 64 |
| HybriDNA-3B | 3B | 16 | 4096 | 8192 | 128 | 8 | 64 |
| HybriDNA-7B | 7B | 32 | 4096 | 8192 | 128 | 8 | 64 |

Pretraining on Multi-Species Genomes

#### Pretraining Corpus

HybriDNA was pretrained on a taxonomically diverse genomic corpus curated from the Nucleotide Transformer dataset^18^ derived from NCBI RefSeq^50^, comprising approximately 160 billion nucleotides from 816 training species spanning bacteria (647 species, 16.5 B bp), fungi (44 species, 2.0 B bp), invertebrates (37 species, 19.9 B bp), protozoa (9 species, 0.45 B bp), mammalian vertebrates (28 species, 65.2 B bp), and non-mammalian vertebrates (51 species, 57.4 B bp), with a held-out validation set of 35 species (13.25 B bp). HybriDNA uses single-nucleotide tokenization (A/C/G/T), preserving resolution for SNPs, point mutations, and exact transcription factor binding site boundaries; the sub-quadratic complexity of the hybrid architecture makes the resulting longer sequences computationally tractable.

To progressively build long-range modeling capacity, we used a curriculum strategy with three context-length stages: (1) 8,192 tokens for 500,000 steps at 0.5 M tokens per batch (∼250 B tokens, ∼1.5 epochs over the corpus); (2) 32,768 tokens for an additional 10,000 steps to capture promoter-proximal regulatory dependencies; and (3) 131,072 tokens for a final 10,000 steps to enable modeling of distal enhancer-promoter interactions and other long-range regulatory architectures. All models were trained with the standard next-token prediction objective using AdamW optimizer^51^ (β_1_ = 0. 9, β_2_ = 0. 95, ϵ = 1*e* − 8, weight decay 0.1), with learning rates of 1e-3, 6e-4, and 1e-4 for the 300M, 3B, and 7B variants respectively, 2,000 warm-up steps, and cosine decay. Training used mixed-precision (BF16), DeepSpeed ZeRO Stage 1 optimization, and Flash-Attention 2^52^.

##### Reverse-Complement Echo Embeddings for Discriminative Tasks

Decoder-only autoregressive models cannot incorporate downstream (3’) information when generating embeddings for upstream (5’) positions. This is particularly limiting for DNA, where regulatory function depends on bidirectional context: transcription factors recognize motifs flanked by sequences on both sides, and many DNA-binding proteins read motifs on either strand. Encoder-only models (e.g., DNABERT-2, Nucleotide Transformer) capture bidirectional context through masked language modeling but sacrifice generative capability. To preserve HybriDNA’s generative capacity while enabling competitive performance on discriminative tasks, we introduce reverse-complement (RC) echo embeddings, inspired by recent advances in text embedding from decoder-only models^53^.

During fine-tuning, the input sequence x of length L is transformed into an “echo” input by concatenation with either a copy of itself (*x*_*echo*_ = [*x*; *x*]) or its reverse complement (*x*_*echo*_ = [*x*; *RC*(*x*)]), where RC(·) denotes A↔T and C↔G substitution followed by sequence reversal. The model processes the echo input autoregressively. When generating hidden states for positions L+1 to 2L, the causal attention mechanism attends back over the complete first half, effectively providing bidirectional context. For classification tasks, we extract a pooled embedding by averaging hidden states from the second half of the echo and feed it through a task-specific linear classification head. The 2× sequence length increase introduced by the echo translates to only ∼1.5× wall-clock time during fine-tuning, because the sub-quadratic Mamba2 layers scale linearly with length and only the periodic Transformer layers incur quadratic cost.

##### Benchmark Evaluation and Fine-Tuning Procedures

To systematically evaluate HybriDNA’s capacity for genomic sequence understanding, we benchmarked its performance on two established evaluation suites: the Genome Understanding Evaluation (GUE) benchmark and the BEND benchmark.

#### Genome Understanding Evaluation (GUE)

GUE^16^ aggregates 28 datasets across 7 task categories, with input sequence lengths ranging from 70 to 512 base pairs. The benchmark serves as a standardized evaluation suite for multi-species genome classification, encompassing the following tasks: (1) Promoter Detection—binary classification of 300-bp sequences as promoter or non-promoter regions in human (3 subtasks: all, notata, tata); (2) Core Promoter Detection—identification of core promoter elements (3 subtasks); (3) Splice Site Prediction—identification of donor and acceptor splice junctions; (4) Transcription Factor Binding Site Prediction—binary classification across 5 human and 5 mouse transcription factors; (5) Epigenetic Marks Prediction—histone modification prediction in yeast; and (6) COVID Variant Classification—viral genome mutation classification.

For GUE fine-tuning, we adopted the experimental settings from DNABERT-2, including task-specific warmup steps and training/validation schedules. All model parameters were fine-tuned (full fine-tuning) using the Adam optimizer with learning rate 5e-5, batch size 32, and weight decay 0.01. For embedding extraction, we evaluated three strategies: (a) the hidden state of the final token for standard fine-tuning, (b) the mean-pooled hidden states from the latter half of the duplicated sequence for repetition echo embedding (denoted with suffix “-E”), and (c) the mean-pooled hidden states from the reverse complement echo (denoted with suffix “-RCE”). A linear classification head was applied on top of the extracted embeddings. We report Matthews Correlation Coefficient (MCC) for all tasks except COVID Variant Classification, for which we report F1 score following the original GUE specification. Results are reported as mean ± standard deviation across three random seeds.

#### BEND Benchmark

BEND^54^ evaluates DNA language models on biologically meaningful tasks derived from functional genomics assays on the human genome. We evaluated three supervised classification tasks: (1) Chromatin Accessibility—predicting DNase-seq and ATAC-seq signals indicating open chromatin regions; (2) Histone Modification—predicting chemical modifications to histone proteins that regulate chromatin structure; and (3) CpG Methylation—predicting DNA methylation patterns at CpG sites.

For BEND fine-tuning, we followed the original benchmark protocol: the pretrained model embeddings were frozen, and only a downstream two-layer convolutional neural network (CNN) was trained for classification. Training proceeded for 100 epochs with learning rate 3e-3 and linear decay to zero. For autoregressive models (HyenaDNA, HybriDNA), we used the mean of hidden states across all sequence positions as the embedding representation. For echo embedding variants, we used the mean of hidden states from the repeated sequence portion. Model selection was based on lowest validation loss, and we report mean AUROC on the held-out test set.

### Computational Efficiency Analysis

To quantify the computational advantages of the hybrid architecture, we compared HybriDNA against a standard Transformer baseline and Striped-Hyena (a hybrid of Transformer and convolutional Hyena operators used in Evo^21^) at a matched scale. All three models comprised approximately 300M parameters and were evaluated on a single NVIDIA A100 GPU (80 GB memory), using Flash Attention 2 for the Transformer and optimized CUDA kernels for the Mamba2 layers. We measured (1) training throughput (tokens/second) with batch size maximized to fit GPU memory, and (2) peak GPU memory consumption (GB) with batch size fixed at 1. HybriDNA achieved substantially higher throughput and lower memory usage than both baselines across all tested context lengths (Figure 1C–1D). The throughput advantage grew with increasing context due to Mamba2’s sub-quadratic complexity: at 49,152 tokens HybriDNA reached up to 4× the throughput of the Transformer baseline, and at 65,536 tokens only HybriDNA could operate within the 80 GB budget; both Transformer and Striped-Hyena baselines encountered out-of-memory errors at shorter contexts.

#### Reinforcement learning benchmark

To evaluate computational efficiency under reinforcement learning workloads, we benchmarked HybriDNA against Evo2 across three model scales using identical PPO settings (512 sequences per iteration, batch size 64, mini-batch size 32, 200-bp generation length) on a single NVIDIA A100 GPU (80 GB). We measured wall-clock seconds per training iteration and peak GPU memory consumption for HybriDNA-300M, HybriDNA-3B, and HybriDNA-7B against Evo2-1B and Evo2-7B. Because Evo2-7B exceeded the 80 GB memory budget at full batch size, we additionally evaluated Evo2-7B at 3/4 of the original batch size; full-batch Evo2-7B is reported as out-of-memory. HybriDNA-7B was the only 7B-scale model that completed RL training under the 80 GB constraint without batch reduction, while HybriDNA-300M and HybriDNA-3B both achieved faster per-iteration speeds than Evo2-1B (Figure 1E–1F).

##### Cell-type-specific CRE Generation

To demonstrate HybriDNA’s utility for functional genomic engineering, we developed a two-stage optimization pipeline for generating synthetic cis-regulatory elements (CREs) with cell-type-specific activity profiles. This pipeline combines supervised fine-tuning (SFT) to establish regulatory sequence generation capability with reinforcement learning (RL) to explicitly optimize for cell-type specificity.

##### Supervised Fine-Tuning on MPRA Data

###### Training Data

We curated cell-type-specific CRE sequences from massively parallel reporter assay (MPRA) datasets available through ENCODE^55^ and the CODA framework^36^. The training corpus comprised experimentally validated regulatory elements from three human cell lines representing distinct lineages:

- **K562:** Chronic myelogenous leukemia cells (erythroid/myeloid lineage)
- **HepG2:** Hepatocellular carcinoma cells (hepatocyte lineage)
- **SK-N-SH:** Neuroblastoma cells (neural lineage)

Detailed sequence selection criteria and source ENCODE accessions are listed in Supplementary Table S1.

###### Data Preprocessing and Cell-Type Specificity Labeling

To identify sequences with strong cell-type-specific activity, we scored all candidate CRE sequences (200 bp each) using the Malinois convolutional neural network from the CODA framework, which predicts regulatory activity across the three cell types. For each cell type, we designated sequences as “high activity” if their Malinois score ranked in the top 25th percentile, and “low activity” if ranked in the bottom 25th percentile. A sequence was classified as cell-type-specific if it exhibited high activity in exactly one cell type and low activity in the remaining two cell types. This stringent filtering criterion ensured that training sequences possessed unambiguous cell-type selectivity rather than broad regulatory activity.

###### Prompt-Conditioned Generation

To enable controllable generation, we introduced cell-type-specific prompt tokens ([K562], [HepG2], [SKNSH]) into HybriDNA’s vocabulary. The embedding layer was expanded to accommodate these new token IDs, which were randomly initialized and learned during fine-tuning. Each training sequence was prepended with its corresponding cell-type label, enabling the model to learn the association between prompt tokens and the regulatory grammar specific to each cell type. The generative objective conditions next-nucleotide prediction on the cell-type prompt:

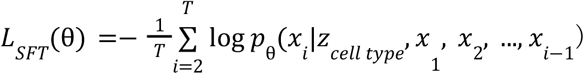

where *z*_*cell type*_ ∈ {[*K*562], [*HepG*2], [*SKNSH*]} is the cell-type prompt token.

###### Fine-Tuning Procedure

The pretrained HybriDNA-300M checkpoint was fine-tuned on the cell-type-specific CRE dataset using a random 80/20 train-validation data split for three epochs. The checkpoint with the lowest validation SFT loss was saved and used in the end. The SFT checkpoint was shared for all target cell types to serve as initialization for subsequent reinforcement learning optimization. SFT was conducted on a single NVIDIA A100 (80 GB) GPU for approximately 5 hours.

### Reinforcement Learning for Cell-Type Specificity

While supervised fine-tuning enables HybriDNA to generate sequences resembling the training distribution, prompt conditioning alone is insufficient to guarantee strong cell-type specificity. Generated sequences may converge toward broadly active but weakly specific regulatory patterns that satisfy the training objective without achieving the desired selectivity. To explicitly optimize for cell-type-specific activity, we implemented a reinforcement learning framework using Proximal Policy Optimization^56^ (PPO) with several domain-specific adaptations.

#### Reward Model

We employed the Malinois convolutional neural network from the CODA framework^36^ as the reward model. Malinois was trained on large-scale MPRA data to predict regulatory activity and takes a 200-bp DNA sequence as input, outputting predicted activity scores {*a*_*K*562_, *a*_*HepG*2_, *a*_*SKNSH*_} for each cell type. To accommodate the model’s expected input length, generated sequences were padded with flanking MPRA vector sequences to match the training context.

#### Reward Function (MinGap Score)

For a sequence generated with target cell type *c*, we define the base reward as the MinGap score with off-target penalty scaling:

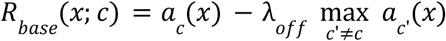

where *a*_*c*_(*x*) is the predicted activity in target cell type *c*, 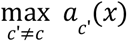 is the maximum predicted activity across non-target cell types, and λ_*off*_ is the off-target penalty coefficient. The off-target penalty coefficient λ_off was linearly ramped from 1.0 at iteration 0 to 5.0 at iteration 1000 to encourage broad exploration in early training and stronger specificity at convergence.

#### Quality-Based Reward Shaping

To accelerate convergence, we applied multiplicative reward shaping based on four reward thresholds. Sequences with *R*_*base*_ ≥ 7 received a 1.5× multiplier; sequences with 5 ≤ *R*_*base*_ < 7 received a 1.3× multiplier; sequences with 2 ≤ *R*_*base*_ < 5 received a 1.1× multiplier; sequences with *R*_*base*_ < 2 were unscaled. Final rewards were clipped to [−2.0, 10.0].

#### Temporal Credit Assignment

Because the Malinois reward model evaluates complete 200-bp sequences, the terminal reward was distributed uniformly across the final 5 generated nucleotide positions, providing a denser learning signal than full-sequence terminal-only assignment.

#### Constrained Sequence Generation

To ensure all generated sequences contained only valid DNA nucleotides, we masked all non-{A,C,G,T} vocabulary logits to negative infinity before nucleus sampling (top-p = 0.9, temperature = 1.0).

#### PPO Implementation

We optimized the SFT-initialized policy using Proximal Policy Optimization with the clipped surrogate objective:

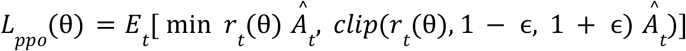

where 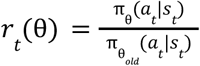 is the probability ratio, 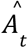 is the advantage estimate computed using Generalized Advantage Estimation (GAE), and ϵ = 0. 2 is the clipping parameter.

The full training objective combined policy, value, entropy, and KL divergence terms:

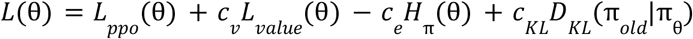

where *c*_*v*_ = 0. 5 is the value loss coefficient, *c*_*e*_ = 0. 05 is the entropy bonus coefficient encouraging exploration, and *c*_*KL*_ = 0. 01 is the KL penalty coefficient maintaining proximity to the reference policy.

#### Value Function

A separate value head was attached to HybriDNA’s final hidden layer, consisting of shared feedforward layers followed by prompt-specific output heads. This architecture allows the value function to learn cell-type-specific baseline estimates while sharing representational capacity across prompts.

#### Experience Replay

To improve sample efficiency and training stability, we maintained an experience replay buffer of 512 trajectories collected from previous PPO iterations. Each trajectory corresponds to one generated 200-bp sequence and stores the full set of (state, action, reward, value, log-probability) tuples required for PPO updates. At each iteration, current on-policy trajectories were combined with replayed trajectories at a 15% replay ratio, sampled with a recency bias (70% drawn from the most recent 25% of the buffer, 30% from older entries).

### Sequence feature analysis

#### ΔK-mer representation of iterative sequences

For each iteration *t*, the sequence set was converted into a k-mer presence ratio vector *x*_*t*_, where the value of each k-mer was defined as the fraction of sequences in that iteration in which the k-mer was observed at least once. The Δk-mer vector was defined as Δ*x*_*t*_ = *x*_*t*_ − *x*_*t*−1_.

#### Motif scanning and motif match scoring

Motif scanning was performed using HOMER (findMotifs.pl with the -find option) with the Vertebrata known motif collection provided by HOMER and default parameters, applied to both 200-bp AI-generated sequences and variable-length STARR-seq sequences. For each motif hit, HOMER reported a motif match score (MotifScore), defined as the log-odds score of the motif matrix and higher scores indicate stronger agreement between the matched sequence and the motif model.

#### Motif matrix definition

For iterative analyses, each iteration consisted of 513 sequences. Motif hits were processed independently for each iteration. Motif match scores below a threshold of 6 were discarded. For each iteration, motif scores were summed across all sequences to generate an iteration-level motif profile. Iteration-level profiles were merged to construct a motif-by-iteration feature matrix, with motifs as rows and iteration indices as columns. Missing values were filled with zero, and columns were ordered according to iteration.

For final generated sequences, motif-based feature matrices were constructed at the sequence level. For each sequence-motif pair, the maximum motif match score was retained. A sequence-by-motif matrix was then generated, with rows representing sequences and columns representing motifs, and missing values filled with zero. For datasets containing more than 10000 sequences, feature matrices were z-score scaled across sequences prior to downstream analysis.

#### Motif-based stage definition

For each iteration, the sequence set was converted into a motif presence ratio vector *m*_*t*_, where the value of each motif was defined as the fraction of sequences in that iteration in which the motif was observed at least once.

Non-negative factorization was performed on the motif presence ratio matrix of each task defined as *M* = [*m*_1_, *m*_2_, …, *m*_1000_] ^*T*^. Three components were used for downstream stage definition based on a combined evaluation of the cophenetic correlation coefficient and residual sum of squares (RSS) across candidate factorization ranks. Component weight trajectories were smoothed using a centered moving average. Stage boundaries were defined as the midpoints between adjacent peaks of the smoothed component trajectories.

### Construction of the shared sequence grammar space

The concatenated Δ6-mer matrix was constructed by pooling the first 40 Δ6-mer vectors from each of the three tasks and was used to compute the covariance matrix, followed by eigendecomposition. The shared eigenvalue spectrum was converted into an effective rank *K* based on the entropy of the normalized eigenvalues. The top *K* covariance eigenvectors defined the shared dynamic basis. Each 6-mer was then represented as

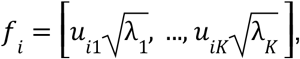

where *u*_*ij*_ denotes the loading of the *i*-th 6-mer on the *j*-th shared dynamic axis, and λ_*j*_ denotes the corresponding eigenvalue.

#### Effective rank estimation

The effective rank *K* of the shared covariance matrix was defined from the entropy of its normalized eigenvalue spectrum. For eigenvalues λ_*i*_, normalized weights were computed as 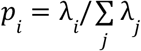, followed by spectral entropy 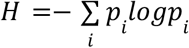. The effective rank was then given by *K* = *ceiling*(*exp*(*H*)), representing the number of covariance dimensions effectively contributing to the shared variance structure.

#### Plasticity score estimation

For each 6-mer, the plasticity was defined as an eigenvalue-weighted loading energy:

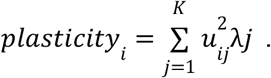

#### Nonlinear embedding by UMAP and visualization

The 6-mer feature vectors were z-score standardized and embedded into two dimensions using UMAP (n_neighbors=30, min_dist=0.15, metric=“euclidean”, random_state=132).

For visualization, plasticity scores were multiplied by a scale factor of 10000. The plasticity and GC fraction of each 6-mer were separately rendered as background landscapes on the 2D grammar space using Gaussian kernel smoothing on a regular grid (GRID_SIZE = 400, SIGMA = 0.04).

#### Projection of iterative trajectories onto the grammar space

For each iteration, the sequence set was converted into a 6-mer presence ratio vector and normalized to sum to 1. The iteration was then projected into the UMAP space as the weighted centroid of the 6-mer coordinates. The shared initial point was defined as the centroid of the task-specific predicted initial points across the three tasks.

### Task-level windowed PCA analysis of sequence update dynamics

#### Calculation of windowed PC contributions

For each task, PCA was performed once on the task-level Δ6-mer matrix to define a shared set of dynamic axes across the full trajectory. Ten principal components were retained for downstream analysis based on the average effective rank estimated from 20-iteration windows. A sliding window of 20 iterations with a step size of 5 iterations was then moved across the trajectory.

Within each window *w*,the root-mean-square (RMS) magnitude of each retained PC score was computed, giving a window-level magnitude vector 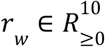. The RMS magnitudes were then smoothed across adjacent windows using a centered rolling mean with window size 2 and normalized to sum to one, yielding the windowed PC contribution vector *c*_*w*_ = (*c*_*w*,1_, …, *c*_*w*,10_), where *c*_*w,j*_ denotes the relative contribution of PC *j* in window *w*.

#### Definition of program transition point

Candidate transition points were identified from changes in the smoothed windowed contribution vectors. For transition detection, only the first five retained PCs were used: the vector 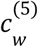 was defined by restricting *c*_*w*_ to PC1-PC5 in the global PCA ordering and renormalizing it to sum to one. A mutation score was then computed for each window as

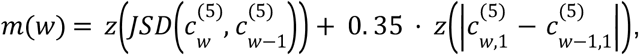

where *z*(·) denotes z-score normalization across windows and JSD denotes Jensen-Shannon divergence.

Candidate peaks were taken as local maxima of *m*(*w*) above the 75% and separated by at least 80 iterations. A candidate peak was retained as program transition point only if it passed an additional stability filter based on the full 10-dimensional contribution vectors:

letting

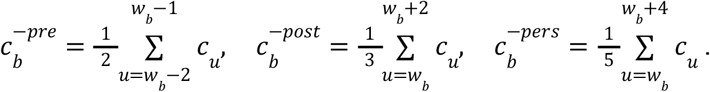

The peak was retained only if

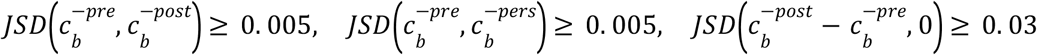

#### Definition of the initial program and transition-associated update programs

In addition to transition-derived programs, an initial program *P*_0_ was defined from the first window by selecting the top three PCs with the largest entries in *c*_0_. For each transition point *b*, a transition-associated update program *P*_*b*_ was defined from the positive contribution change across the transition,

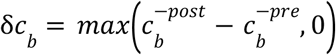

The three PCs with the largest entries in δ*c*_*b*_ were selected as the sparse signature of *P*_*b*_.

Their normalized values were used as program weights *a*_*b,j*_, such that 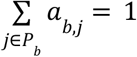. If δ*c*_*b*_ contained no positive entries, the signature was instead taken from the top three PCs in 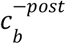.

#### Relative activity of initial and transition-associated programs

For each program *P*_*b*_, a raw window-level score was first computed as the weighted sum of PC contributions,

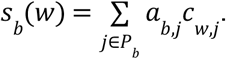

To reflect delayed onset after a transition, transition-associated programs were multiplied by a smooth activation gate defined from the window-center iteration relative to the corresponding transition point, using a clipped smooth-step function over a transition scale of 24 iterations; the initial program *P*_0_ used a constant gate of 1. The gated score was then rectified to nonnegative values, transformed by a power of 1.85, and smoothed on the active segment using a Savitzky–Golay filter (target window length 141, polynomial order 3). Finally, the activities of all programs were renormalized within each window to sum to one, yielding the relative program activity

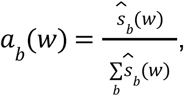

where 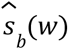 denotes the gated and smoothed nonnegative score for program *b*.

#### Projection of initial and transition-associated programs into 6-mer space

For each program *P*_*b*_, a sparse program vector *g*_*b*_ ∈ *R*^10^ was first defined in PC space, with *g*_*b,j*_ = *a*_*b,j*_ for *j* ∈ *P*_*b*_ and *g*_*b,j*_ = 0 otherwise. Let *L* ∈ *R*^10×4096^ denote the PCA loading matrix learned from the standardized Δ6-mer matrix. The program-specific 6-mer tendency was then obtained by direct projection of the sparse program vector onto the 6-mer loadings as

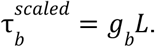

Empirical two-sided p-values, Benjamini–Hochberg adjusted q-values, and rank statistics were calculated across all 6-mers using 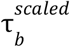 as the primary tendency score.

#### Projection of initial and transition-associated programs into motif space

Each motif PWM was first converted into a strand-collapsed 6-mer probability distribution. For a motif *m*, probabilities were averaged across all overlapping 6-bp windows of the PWM. Forward and reverse-complement 6-mer distributions were then merged by elementwise maximum and renormalized, yielding a motif-specific 6-mer distribution *d*_*m*_ = (*d*_*m*,1_, …, *d*_*m*,4096_). The value of motif tendency of motif *m* for program was defined as the expectation of the program’s 6-mer tendency under this motif-induced distribution,

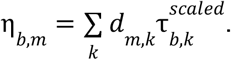

Here, the sign of η_*b,m*_ indicates whether the motif aligns more strongly with the positive or negative side of the program tendency axis.

To assign a direction more directly related to post-transition motif dynamics, motif tendency was further direction-corrected using the observed post-iteration motif change. For each program transition point *b*, the motif dynamic of post 10-iteration was used to determine the final direction. If 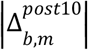 exceeded the threshold defined as the median absolute post-iteration change across the motif set, the motif tendency score was reoriented by the sign of the observed post-branch change,

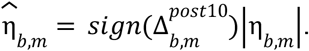

Motifs with 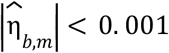 were discarded, and empirical two-sided p-values, Benjamini–Hochberg adjusted q-values, and rank statistics were calculated from 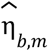.

### Motif-based clustering and heatmap visualization in AI-generated sequences

Motif feature matrices were constructed as described above and used as input for dimensionality reduction and clustering analyses. Low-dimensional representations were computed using UMAP as implemented in the uwot package. A fixed random seed (123) was applied to ensure reproducibility. UMAP was performed with PCA initialization (init = “pca”) and approximate nearest-neighbor search using Annoy (nn_method = “annoy”). Dataset-specific parameters were applied to accommodate differences in motif space density and sample size. For HepG2-specific sequences, UMAP was performed with n_neighbors = 40, min_dist = 0.05, and n_epochs = 800; for K562-specific, n_neighbors = 40, min_dist = 0.01, and n_epochs = 1000; for SK-N-SH, n_neighbors = 60, min_dist = 0.05, and n_epochs = 800. All embeddings were generated in two dimensions.

Clustering was performed on the resulting UMAP coordinates using HDBSCAN (https://cran.r-project.org/web/packages/dbscan/vignettes/hdbscan.html) to identify density-based groups while allowing noise points. The minPts parameter was adjusted per dataset (HepG2: 80; K562: 30; SK-N-SH: 110) to reflect differences in local point density. Points labeled as noise were excluded from downstream analyses, and scatter plots were generated using ggplot2 (https://ggplot2.tidyverse.org/).

To visualize motif usage patterns across clusters, heatmaps were generated from the original motif feature matrix. Motif scores were joined with cluster assignments, and only non-noise samples were retained. The feature matrix was transposed to motif-by-sample format, and motifs present in fewer than 600 samples were filtered out. Cluster annotations derived from HDBSCAN labels were added as column annotations. Heatmaps were rendered using ComplexHeatmap (https://bioconductor.org/packages/release/bioc/html/ComplexHeatmap.html), with cluster identity indicated by a categorical color scale.

### Compare with endogenous CREs

Endogenous CREs were obtained from reproducible HepG2 and K562 STARR-seq peaks processed by Lee et al. using datasets ENCFF047LDJ and ENCFF045TVA, respectively. For each cell type, endogenous CREs and AI-generated candidate CREs were represented using motif-based sequence feature matrices as described above and analyzed jointly.

#### Dimensionality reduction by UMAP

Dimensionality reduction was performed by applying PCA to the combined feature matrix, retaining the first 30 principal components. Two-dimensional embeddings were generated using UMAP on the PCA transformed features with the following parameters:metric = “cosine”, init = “spectral”, nn_method = “annoy”, n_components = 2, n_epochs = 1000, random seed 213 and n_neighbors = 30 for HepG2 and 50 for K562.

#### Two-step clustering and motif heatmap visualization

Two-step clustering was performed on the two-dimensional UMAP embeddings. First, K-means clustering was applied with 20000 centers (nstart = 80, iter.max = 100) to partition the embedding space. Second, HDBSCAN was run on the K-means cluster centers (minPts = 50 for HepG2 and 70 for K562) to identify dense regions. The resulting HDBSCAN cluster assignments were then mapped back to the motif-based feature matrices.

For each non-noise cluster, up to 60 endogenous sequences were randomly sampled with replacement using a fixed random seed (set.seed = 668), and all AI-generated sequences were retained. The motif-based feature matrix was processed using Seurat v5.3.0 to identify the 300 most variable motifs (selection.method = “vst”). Motifs present in fewer than 200 sequences were excluded and visualized as heatmap using the ComplexHeatmap.

#### ChromBPNet prediction

Chromatin accessibility of 200-bp candidate CREs was evaluated using ChromBPNet with pretrained ATAC-seq models for HepG2 (ENCFF137WCM) and K562 cells (ENCFF984RAF). For ChromBPNet prediction, each 200-bp candidate sequence was embedded at a fixed position within a standardized 2,114 bp input sequence. Specifically, candidate sequences replaced the 200-bp segment corresponding to chr16: 28,930,800–28,930,999 within a fixed 2,114 bp genomic window spanning chr16:28,929,800–28,931,913.

Predictions were generated using five independently trained no-bias ChromBPNet models per cell line, with averaged outputs used as the final chromatin accessibility profiles. For each sequence, the mean predicted accessibility within the insertion window (positions 470–570 of the model output) was extracted and used to represent the chromatin accessibility of the corresponding CRE.

### Motif phrase analysis

Motif syntax analysis was performed for three selected motifs. Candidate CREs containing all three motifs were extracted, and motif positions were represented by the offset reported by HOMER. For each analysis, one motif was designated as the reference, and each occurrence of it was treated as the coordinate origin. The relative positions of the other two motifs with respect to this origin were calculated, forming a set of three-motif position triplets. The resulting two-dimensional distributions of motif triplets were visualized as 3D bar plots using persp in R.

### MPRA library design

#### Negative control

Negative control sequences were derived from GM12878 ATAC-seq open chromatin regions. Multiple GM12878 ATAC-seq BED files were first intersected to identify regions consistently accessible across datasets, and an aggregated ATAC-seq score (sum_score) was calculated for each region by summing the peak scores across all GM12878 ATAC-seq datasets. Candidate regions were then stringently filtered by excluding any intervals overlapping ATAC-seq or DNase-seq peaks from K562, HepG2, or SK-N-SH, and regions shorter than 200 bp were removed. The remaining regions were ranked in descending order first by sum_score per base pair and then by sum_score. From the top 100 ranked regions, 50 were randomly sampled (random_state = 42). In addition, 50 regions were randomly sampled from an intermediate ranking range spanning ranks 1001 to 5000 (random_state = 42). These two subsets were combined, and each interval was subsequently partitioned into multiple overlapping 200 bp fragments using an adaptive tiling strategy. The number of fragments was determined by the interval length, and fragment start positions were evenly spaced to ensure complete coverage of the original interval while maintaining a fixed fragment length. All sequences were extracted from the hg38 reference genome. Sequences containing ambiguous bases were excluded. Genomic specificity was assessed using the UCSC Genome Browser BLAT search. Sequences were retained only if they showed a single full-length 200 bp perfect match in the human genome, with no additional alignments longer than 30 bp.

#### Library for cell-type-specific CREs validation

This library comprised 1000 constructs, including 50 negative control sequences, 450 AI-generated candidate CREs (150 per cell type), and reverse complements of all sequences. The 50 negative control sequences were randomly sampled without replacement from the negative control library using a fixed random seed (431). AI-generated CREs were selected based on predicted regulatory activity scores generated by the Malinois model as described above. Sequences were first grouped according to their UMAP-derived clusters. Within each cluster, sequences were ranked using a combined criterion that jointly considered i) the predicted activity score in the target cell type and ii) a cell type specificity score, defined as the difference between the predicted activity in the target cell type and the sum of predicted activities in the two non-target cell types. The combined rank was computed as the sum of the individual ranks for these two metrics, and the top 30 sequences were selected from each cluster. Clusters were then compared based on the mean predicted activity score and mean cell-type–specificity score of their selected sequences. A cluster-level combined rank was derived from these two mean values, and the top five non-noise clusters were retained, yielding a total of 150 AI-generated CREs per cell type.

#### Library for comparison with endogenous

The MPRA library comprised 1,000 constructs, including 400 HepG2 endogenous CREs, 400 AI-generated HepG2-specific candidate CREs, and 200 negative control sequences which were randomly sampled without replacement from the negative control library using a fixed random seed (431). Endogenous CREs identified from HepG2 STARR-seq data were obtained from reproducible HepG2 STARR-seq peaks reported by Lee et al. Peaks shorter than 200 bp were excluded, and each remaining peak was converted into fixed-length 200 bp fragments by generating up to 100 overlapping windows with uniformly spaced start positions to ensure complete coverage of the original interval. These fragments were first scored using boda2, and the top 5000 fragments with the highest MinGap for HepG2 as the target cell type were retained. The retained fragments were subsequently ranked by chromatin accessibility predicted by ChromBPNet using the mean predicted value across the prediction region. Fragments were selected under the additional constraint that selected fragments were separated by at least 20 fragment indices within the same peak. Based on these criteria, the top 400 fragments were retained as representative HepG2 endogenous CREs. AI-generated HepG2-specific candidate CREs were filtered in two steps. The top 1,000 sequences were first selected based on MinGap scores predicted by the boda2 framework using the Malinois model. These sequences were subsequently ranked by chromatin accessibility predicted by ChromBPNet, and the top 400 CREs were retained.

### LentiMPRA analysis

Raw MPRA sequencing reads were first adapter-trimmed using cutadapt v5.1 and merged using PEAR v0.9.6 to generate contiguous sequences for alignment. For libraries containing reverse complement (RC) sequences, separate Bowtie2 v2.5.4 indices were built for the forward and RC sequences. Merged reads were aligned independently to both indices, and only forward-aligned reads were retained to generate counts represented forward and RC sequences respectively. For libraries without RC sequences, a single Bowtie2 index was built and reads were aligned once to extract counts. BAM files were sorted and indexed using samtools v1.22.1, and read counts for each sequence were tabulated. For each sequence, activity scores were computed as a weighted sum of binned signal counts. Specifically, four bins (bin1–bin4) were combined as:

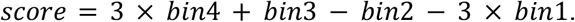

The resulting score was then linearly rescaled to the range [-1,1] using a min–max normalization function, yielding the normalized activity score. For experiments with multiple replicates, normalized scores were averaged across replicates to obtain the final activity score for each sequence.

### Amplification of custom oligonucleotide pools

The oligonucleotide library, synthesized by Twist Bioscience, was cloned into the lentiviral MPRA vector following PCR amplification. Forward (Oligo-F: 5′–AACCGGTGGCCACTTCAT TT–3′) and reverse (Oligo-R: 5′–CCCTCTAGTGACCTTGACAGC–3′) primers, complementary to the predefined 5′ and 3′ adaptor sequences of the oligonucleotide pool, were used for amplification. PCR amplification was performed using Q5 High-Fidelity 2× Master Mix (New England Biolabs, M0492S) under the following conditions: initial denaturation at 98°C for 30 s, followed by 13 cycles of 98°C for 10 s, 58°C for 10 s, and 72°C for 20 s, with a final extension at 72°C for 2 min. The amplified products were resolved on a 1.5% agarose gel, and the fragment of the expected length was extracted and purified using MagPure Gel Extraction Kit (Magen, D2110) for ligation into the linearized lentiviral backbone.

### Lentiviral MPRA library construction

The lentiviral MPRA backbone was based on plasmid pLS-SceI (Addgene #137725)^57^. This vector was modified to introduce an hPGK promoter-driven mCherry cassette downstream of the original reporter and to replace the SceI cloning site with dual BfuAI restriction sites. The modified vector was linearized using BfuAI (New England Biolabs, R0701S) in NEBuffer™ r3.1. The linearized product was separated on a 1% agarose gel, and the backbone fragment of the correct size was gel-extracted and purified using MagPure Gel Extraction Kit (Magen, D2110).

The final MPRA vector contains two expression cassettes: 1) an upstream reporter cassette, in which oligonucleotide libraries are inserted between flanking adaptor sequences within a minimal promoter-driven transcription unit. This unit consists of the minimal promoter, the oligo cloning site, an eGFP sequence fused to a PEST destabilization domain, and a bovine growth hormone polyadenylation signal (bGHpA); 2) a downstream constitutive expression cassette consisting of the hPGK promoter driving mCherry, which serves as a transduction and normalization control. Purified, double-stranded oligonucleotide pool PCR products (250 bp) were cloned into the linearized backbone using a 2× Seamless Cloning Mix (Biomed, CL117-01). For cloning, six independent 20 μL ligation reactions were assembled, each containing 330 ng of linearized backbone and 50 ng of insert DNA. Reactions were incubated at 50°C for 60 min, pooled, and co-precipitated with 4 μg of glycogen (Roche, 10901393001) using isopropanol. The pelleted DNA was resuspended in 5 μL of nuclease-free water. Two separate transformations were performed using Endura™ ElectroCompetent Cells (Biosearch Technologies, 60242), each with 2.5 μL of the purified ligation product. Following transformation, cells were recovered in 2 mL of SOB medium at 37°C for 1 h. The two cultures were combined and plated across forty 15-cm LB agar plates. After overnight growth, colonies were harvested, and plasmid DNA was extracted using a QIAGEN Plasmid Maxi Kit (QIAGEN, 12362).

### Plasmid library quality control

The plasmid library was deep-sequenced for quality control. We designed sequencing primers flanking the oligo insert region, compatible with the Illumina TruSeq system and containing unique dual indices (UDIs) to enable multiplexing (see oligonucleotides).

A two-step PCR strategy was employed. In the first step, the plasmid library was amplified for 10 cycles using the TruSeq adapter primers under conditions otherwise identical to those described for oligonucleotide pool amplification. The PCR product was purified. In the second step, the purified product was used as a template for an 8-cycle amplification with the UDI-indexed primers. The final PCR product was purified and subjected to paired-end sequencing on an Illumina NovaSeq 6000 platform at Annoroad (Beijing, China) to assess library complexity and sequence representation.

### Cell culture

HepG2, K-562, and HEK293T cell lines were kindly provided by Dr. Ting Han’s laboratory, while SK-N-SH (ATCC HTB-11) cells were purchased from the American Type Culture Collection (ATCC). HepG2, K-562, and HEK293T cells were cultured in Dulbecco’s Modified Eagle Medium (Gibco, 11965092). SK-N-SH cells were maintained in Minimum Essential Medium (Gibco, 11095080). All media were supplemented with 10% fetal bovine serum (GeminiBio, 900-108-500) and 1% penicillin–streptomycin (Gibco, 15140122). All cells were maintained at 37 °C in a humidified incubator with 5% CO_2_.

### Lentiviral vector production

293T cells were seeded in 10 cm dishes and grown to approximately 70% confluence prior to transfection. Lentiviral vectors were produced using a second-generation packaging system consisting of psPAX2 (Addgene, 12260) and pMD2.G (Addgene, 12259). For transient transfection, 10 µg of plasmid library, 7.5 µg of psPAX2, and 2.5 µg of pMD2.G were co-transfected into 293T cells using Lipofectamine 3000 (Thermo Fisher Scientific, L3000075) according to the manufacturer’s instructions. Eight hours after transfection, the culture medium was replaced with fresh complete medium. Viral supernatants were collected 48 hours post-transfection, clarified by filtration through a 0.45 µm filter, and used for downstream transduction.

### Lentiviral transduction

For suspension cells, K-562 cells (1 × 10^6^) were resuspended in 10 mL of filtered lentiviral supernatant supplemented with 10 μg/mL polybrene (Yeasen, 40804ES). Cells were subjected to spinfection by centrifugation at 1,500 × g for 1 h at room temperature, followed by incubation at 37 °C for 12 h. For adherent cell lines, HepG2 and SK-N-SH cells were seeded in 6-well plates one day prior to transduction to reach approximately 70% confluence at the time of infection, with 5 mL of filtered lentiviral supernatant added per well. Polybrene was added at a final concentration of 8 μg/mL for HepG2 cells and 5 μg/mL for SK-N-SH cells. HepG2 cells were subjected to spinfection at 1,000 × g for 30 min, whereas SK-N-SH cells were spinfected at 500 × g for 1 h. All cells were incubated with lentiviral supernatant for 12 h post-infection, after which the medium was replaced with fresh complete culture medium. K-562 and HepG2 cells were passaged and expanded at 48 h post-infection. For all lentiviral MPRA experiments, the multiplicity of infection (MOI) was maintained below 0.3 to ensure predominantly single-copy integration. Transduced populations were enriched by flow cytometric sorting at 96 h after infection.

### Flow cytometric sorting and data analysis

Post-transduction cells were washed with DPBS (Thermo Fisher Scientific, 14190144). Adherent HepG2 and SK-N-SH cells were detached using Trypsin-EDTA (Thermo Fisher Scientific, 25200114) at 37 °C (HepG2: 5 min; SK-N-SH: 2 min). Digestion was terminated by adding complete culture medium, and cells were collected by centrifugation and resuspended in MACS buffer (DPBS supplemented with 2% FBS and 2 mM EDTA). Cells were sorted on a BD FACSAria™ III Cell Sorter (BD Biosciences). Untransduced cells were included in each experiment as gating controls. The sorting strategy was as follows: debris was excluded based on FSC/SSC parameters, DAPI-negative cells were gated as viable cells, singlets were selected, and mCherry-positive cells were gated. Within the mCherry-positive population, cells were further subdivided into four equal bins according to the eGFP/mCherry fluorescence ratio. Approximately 1 × 10^6^ cells were sorted and collected from each bin for downstream Illumina sequencing. Flow cytometry data were analyzed using FlowJo software (v10.8.1, BD Biosciences).

### Sequencing library preparation from sorted MPRA cell populations

Genomic DNA from each of the four sorted bins was extracted using the QIAamp DNA Mini Kit (QIAGEN, 51304) following the manufacturer’s instructions.

A two-step PCR was performed using primer sets identical to those used in plasmid library quality control. In the first PCR, 2 µg genomic DNA per 50 µL reaction was amplified with TruSeq adapter primers for 20 cycles. Products were purified using NovoNGS® DNA Clean Beads (Novoprotein, N240), and fragments of 200–300 bp were selected using a 0.9× bead ratio. In the second PCR, purified first-round products served as templates (2 ng per 50 µL reaction) and were amplified with UDI-indexed primers for 10 cycles. PCR products were size selected on a 1% agarose gel, and correctly sized bands were excised and purified using the MagPure Gel Extraction Kit (Magen, D2110). Final libraries were subjected to paired-end sequencing on an Illumina NovaSeq 6000 platform at Annoroad (Beijing, China) to assess sequence distribution.

---

**Supplementary Figure 1.**
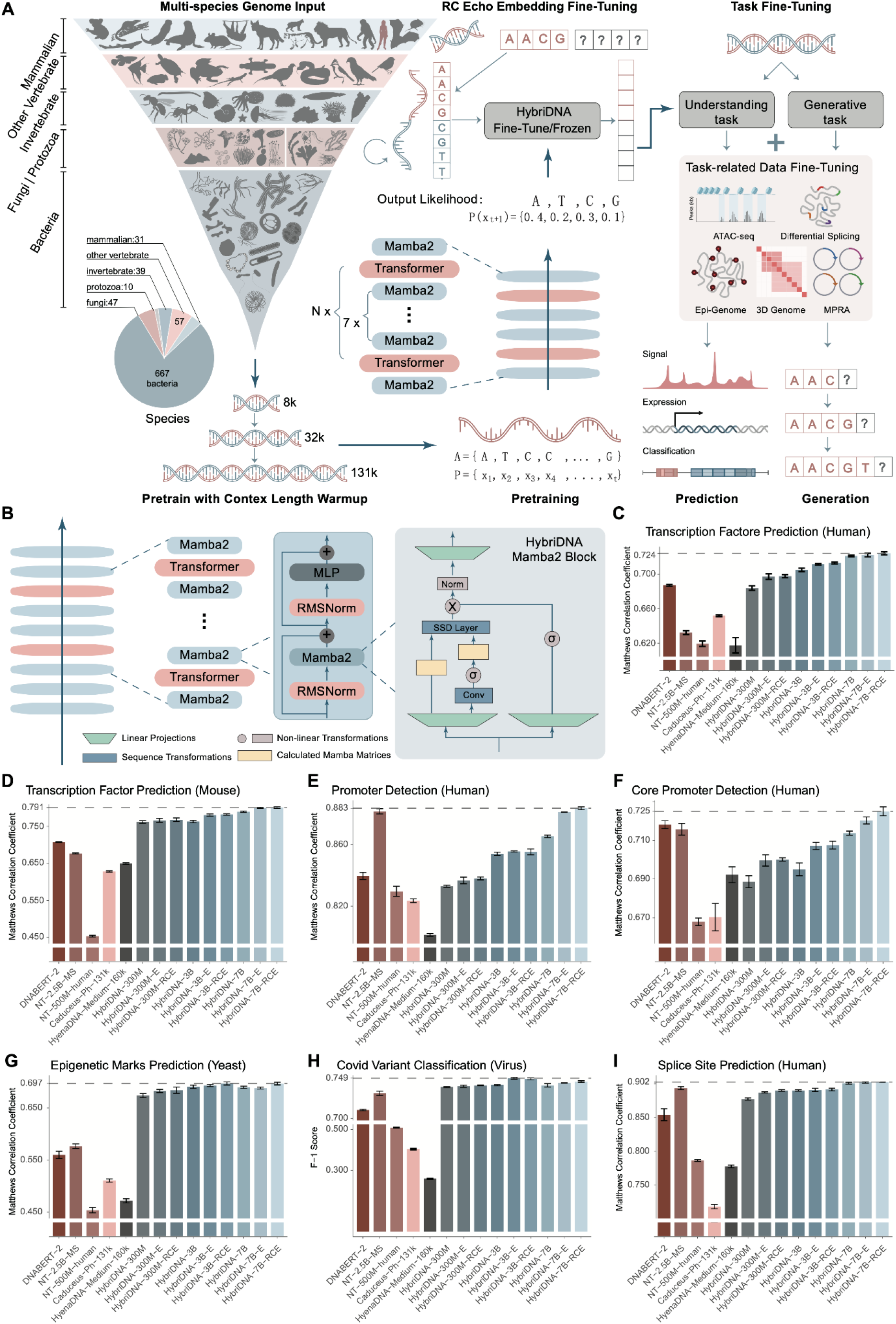
Optimized HybriDNA architecture and benchmarking on genome understanding tasks, related to Figure 1. (A) Schematic of the HybriDNA framework, including multi-species pretraining with context-length warmup, reverse-complement echo fine-tuning, and adaptation to predictive and generative tasks. (B) Detailed architecture of the HybriDNA Mamba2 block. (C–I) Performance of genomic language models on Genome Understanding Evaluation (GUE) tasks. (C) Transcription factor prediction in human. (D) Transcription factor prediction in mouse. (E) Promoter detection in human. (F) Core promoter detection in human. (G) Epigenetic mark prediction in yeast. (H) COVID variant classification. (I) Splice-site prediction in human. Models compared include DNABERT-2, NT-2.5B-MS, NT-500M-human, Caduceus-Ph-131k, HyenaDNA-Medium-160k, and HybriDNA variants. Matthews correlation coefficient (MCC) was used in (C–G) and (I), and F1 score was used in (H). E, Echo Embedding; RCE, Reverse Complement Echo Embedding. Error Bar represents S.D. of three independent runs with random seeds.

**Supplementary Figure 2.**
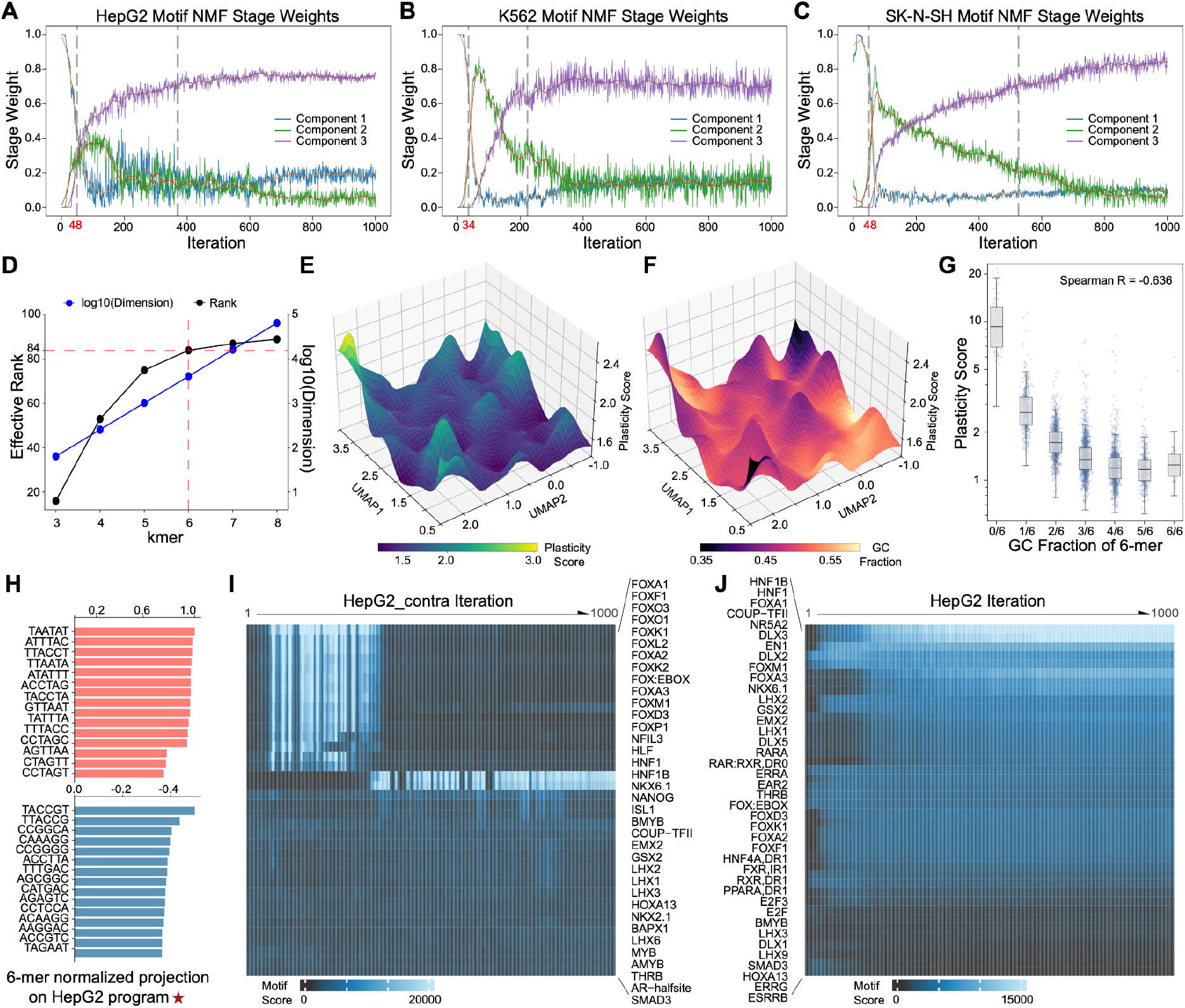
Phase dynamics and optimization trajectories in reconstructed sequence-grammar space, related to Figure 2. (A–C) Non-negative matrix factorization (NMF) of summed motif match scores across 1,000 optimization iterations for HepG2 (A), K562 (B), and SK-N-SH (C). Three latent components are shown. Gray dashed lines indicate putative phase boundaries, defined as the midpoints between adjacent smoothed component peaks. (D) Effect of increasing k-mer length on the dimensionality of sequence-grammar space. Black line indicates effective rank, and blue line indicates the dimensionality of sequence k-mer distributions. (E–F) Gaussian-smoothed 6-mer grammar landscape colored by plasticity score (E) or GC fraction (F). (G) Relationship between plasticity score and GC fraction across all 6-mers. Each point represents one 6-mer (Spearman’s ρ = −0.636). (H) Top positively and negatively weighted 6-mers associated with the first major HepG2 update program at iteration 100. (I-J) Heatmap of motif match scores across HepG2 contra (I) and HepG2 (J) RL iterations. Columns represent iterations and rows represent transcription factor motifs. For panels A–C, I–J, the x axis indicates RL iteration (1–1,000), increasing from left to right; y axes indicate the corresponding plotted features.

**Supplementary Figure 3.**
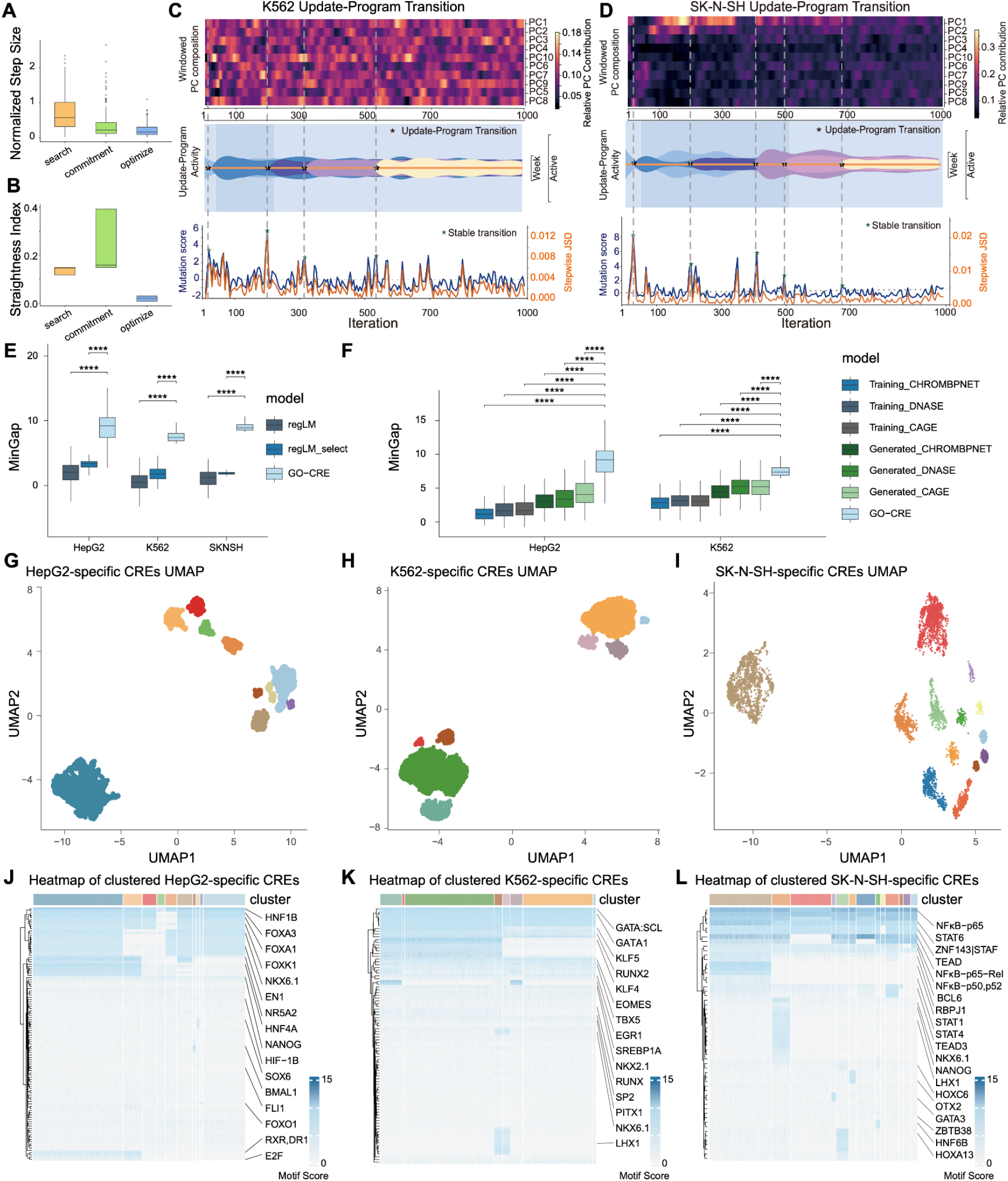
Dissection of generated CREs for iterations and cell-type-specificity, related to Figure 3. (A–B) Trajectory statistics across three representative phases: search, commitment, and optimization. (A) Step size normalized to iteration 2. (B) Straightness index, defined as net displacement divided by total path length within each phase. (C–D) Update-program transitions during K562 (C) and SK-N-SH (D) CRE generation process, including windowed principal component composition, relative principal component contributions, update-program activity, mutation score (blue), and stepwise Jensen–Shannon divergence (orange). Stars indicate emergence of stable transition points. (E) Comparison of MinGap scores of GO-CRE-generated CREs, regLM-generated CREs and 100 selected regLM-generated CREs across HepG2, K562, and SK-N-SH. Statistical significance was assessed using two-sided Wilcoxon rank-sum tests. GO-CRE was compared to each control group within each cell type. **** *p* < 0.0001. (F) Comparison of MinGap scores among GO-CRE generated CREs, DNA-Diffusion generated CREs, and DNA-Diffusion training CREs in HepG2 and K562. For DNADiffusion-generated and training CREs, the top 200 sequences were selected based on ChromBPNet, DNase, or CAGE scores. Statistical significance was assessed using two-sided Wilcoxon rank-sum tests. GO-CRE was compared to each control group within each cell type. **** *p* < 0.0001. (G–I) Individual UMAP visualization of 9,000 post-evolved generated CREs from HepG2 (G), K562 (H), and SK-N-SH (I). (J–L) Heatmap of motif scores across representative transcription factor motifs in post-evolved CREs from HepG2 (J), K562 (K), and SK-N-SH (L). Columns represent individual CREs ordered by hierarchical clustering, excluding the noise cluster (n =7,898, 8,778 and 7,042 for HepG2, K562 and SK-N-SH, respectively). Rows correspond to distinct motifs (n =107, 91 and 55 for HepG2, K562 and SK-N-SH, respectively). Top annotation bars represent color code matched subclusters from G–I.

**Supplementary Figure 4.**
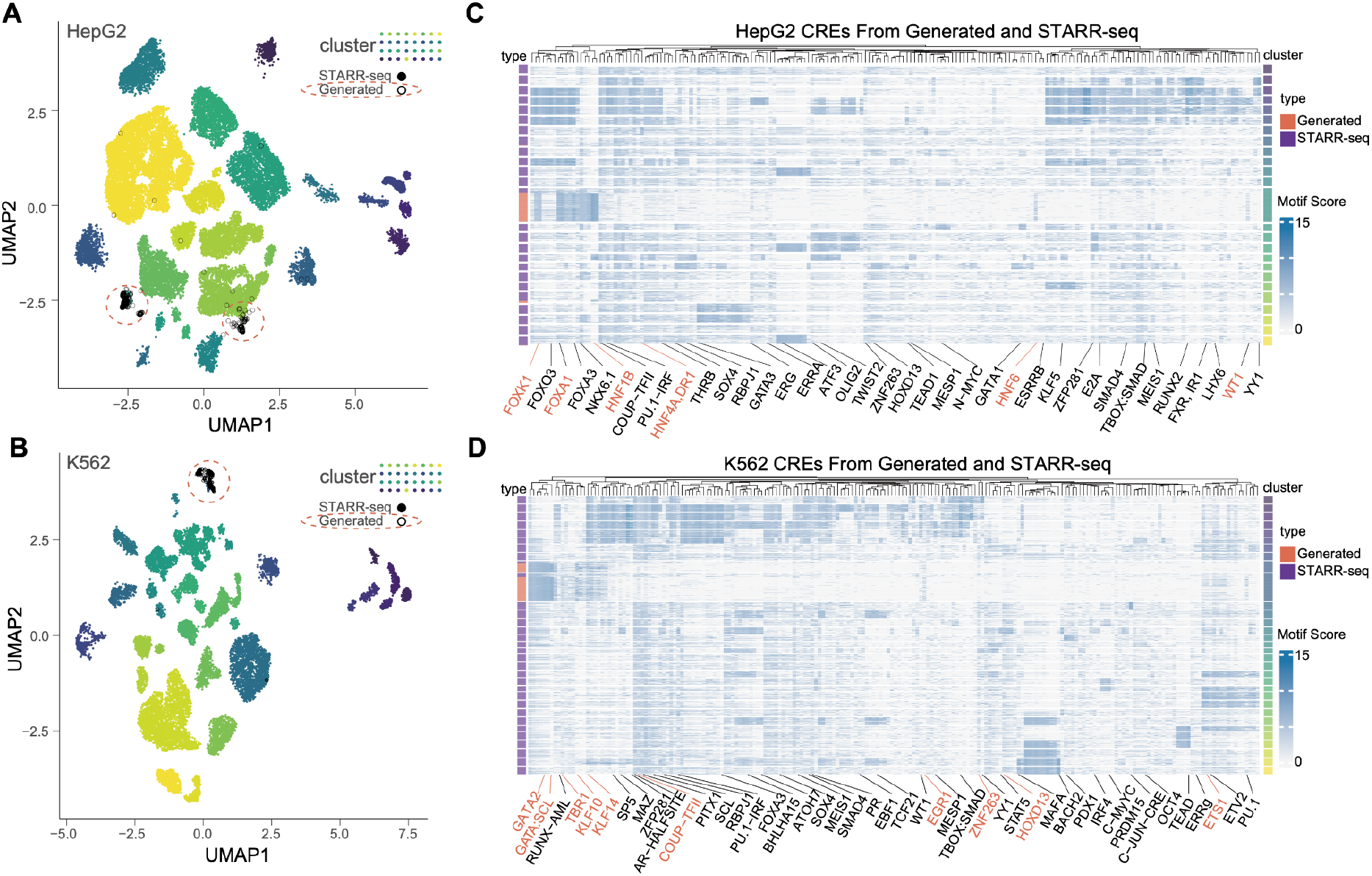
Comparison of generated and endogenous CREs, related to Figure 4. (A–B) Generated CREs projected onto the endogenous CRE UMAP space in HepG2 (A) and K562 (B) cells. Endogenous CREs defined by STARR-Seq are clustered into multiple subtypes indicated in the plots. Projected generated CREs are marked as blank circles. (C–D) Heatmap of motif scores across representative transcription factor motifs in post-evolved CREs from HepG2 (C) and K562 (D). Rows represent individual CREs ordered by hierarchical clustering. Left-side annotation bars represent CRE origin from endogenous (purple) or generated (red). Right-side annotation bars represent color-coded subclusters matched to A–B.

**Supplementary Figure 5.**
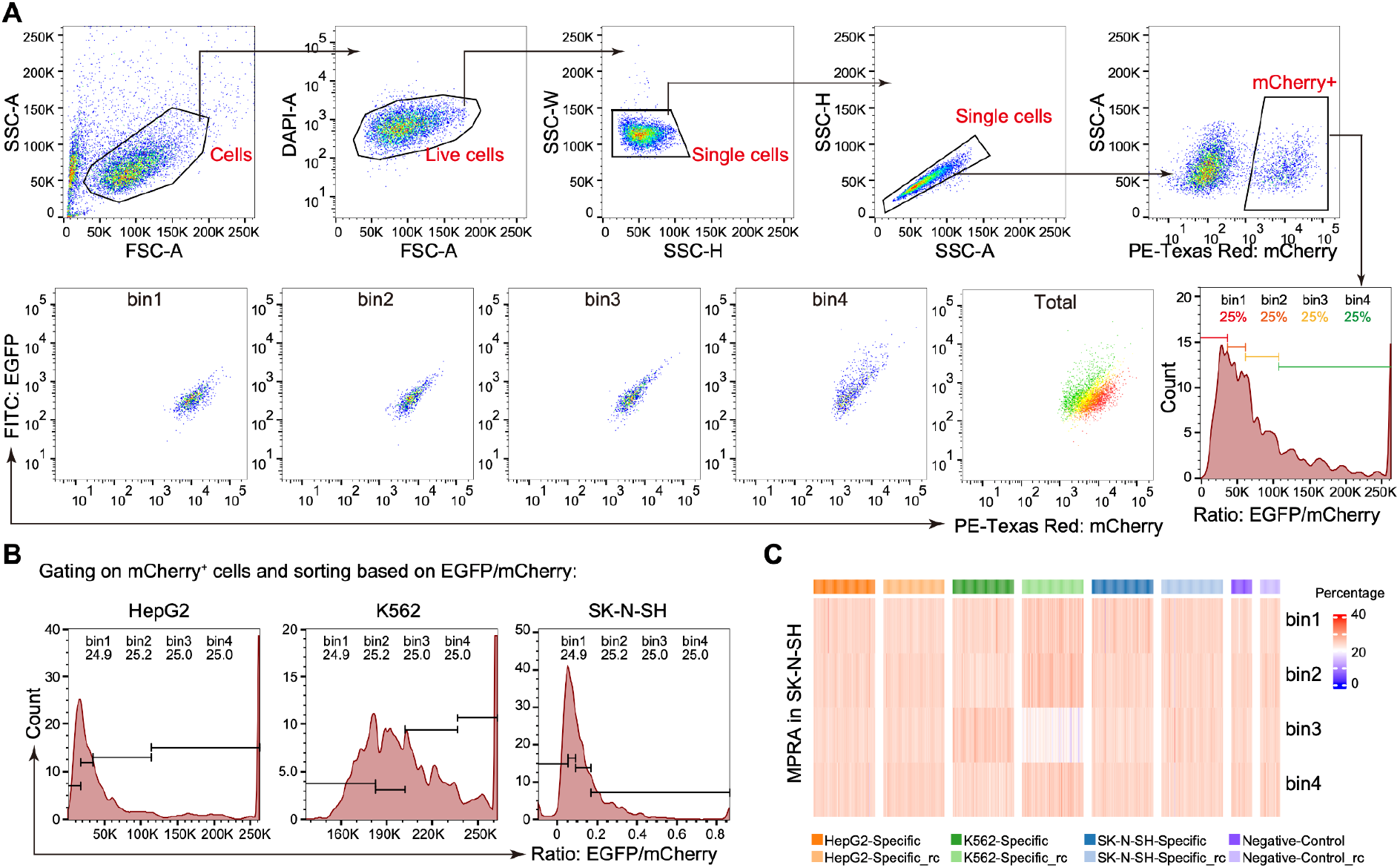
LentiMPRA flow cytometric sort gating strategy and SK-N-SH LentiMPRA profiling, related to Figure 5. (A) Representative gating strategy for sorting LentiMPRA reporter cells. Cells were gated sequentially on intact events, DAPI-negative live cells, singlets, and mCherry-positive cells. The mCherry-positive population was then divided into four bins according to the EGFP/mCherry fluorescence ratio, with bins 1–4 containing approximately equal fractions of cells and corresponding to increasing enhancer activity. (B) Representative histograms of EGFP/mCherry fluorescence ratios in mCherry-positive HepG2, K562, and SK-N-SH cells. Bin boundaries used for sorting and the fraction of cells in each bin are indicated. (C) Heatmap of CRE enrichment across bins 1–4 in SK-N-SH cells. Columns represent individual CREs, and rows represent sorted bins of increasing EGFP/mCherry ratio as defined in (A). Top annotation bars indicate CRE class. RC, reverse-complemented sequence. Data represent 2 independent screens.

**Supplement Table 1:** MPRA datasets for fine-tuning.

| Biosample | Accession |
| --- | --- |
| HepG2 | ENCSR045VPN, ENCSR167HAR, ENCSR222QJT, ENCSR354FXV, ENCSR447RLV, ENCSR463DWN, ENCSR554UKK, ENCSR694VUM, ENCSR712VZT, ENCSR736XUK, ENCSR811QES, ENCSR833BYO, ENCSR843APF, ENCSR847SNQ, ENCSR979TSK, ENCSR632EPR, ENCSR208DGH, ENCSR839LDC, ENCSR215YXA, ENCSR463IRX, ENCSR405QCT, ENCSR255GVC, ENCSR049RIU, ENCSR022GQD, ENCSR509EUX, ENCSR202CHZ, ENCSR787MOJ, ENCSR002UCW, ENCSR760GMX, ENCSR080JVM, ENCSR478ESZ, ENCSR881ERG, ENCSR661ZUO, ENCSR069RCF |
| K562 | ENCSR003TJR, ENCSR135MRI, ENCSR274WZF, ENCSR326HQU, ENCSR341GVP, ENCSR363XER, ENCSR653LWA, ENCSR697PNZ, ENCSR753ZRI, ENCSR820RPI, ENCSR857JFG, ENCSR909JVP, ENCSR971BEX, ENCSR971PLA, ENCSR997GVD, ENCSR460LZI, ENCSR382BVV, ENCSR203UFY |
| SK-N-SH | ENCSR087VPA, ENCSR386XIB, ENCSR505LQA, ENCSR541FWB, ENCSR562GGR, ENCSR563TTU, ENCSR574GPT, ENCSR579QGV, ENCSR623DJZ, ENCSR742AAP, ENCSR833GXJ, ENCSR856SQG, ENCSR959NUX |

**Supplement Table 2:**
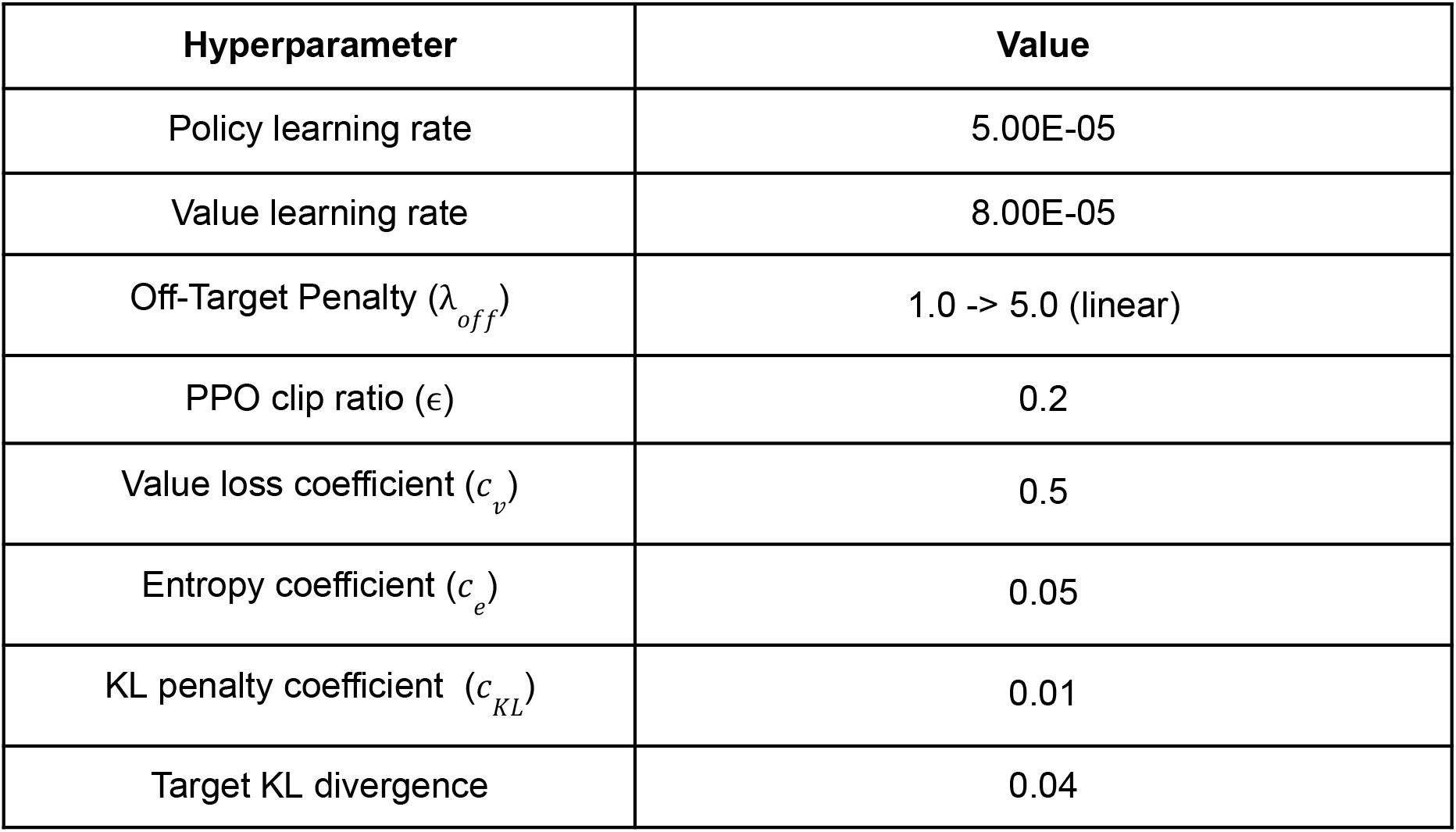
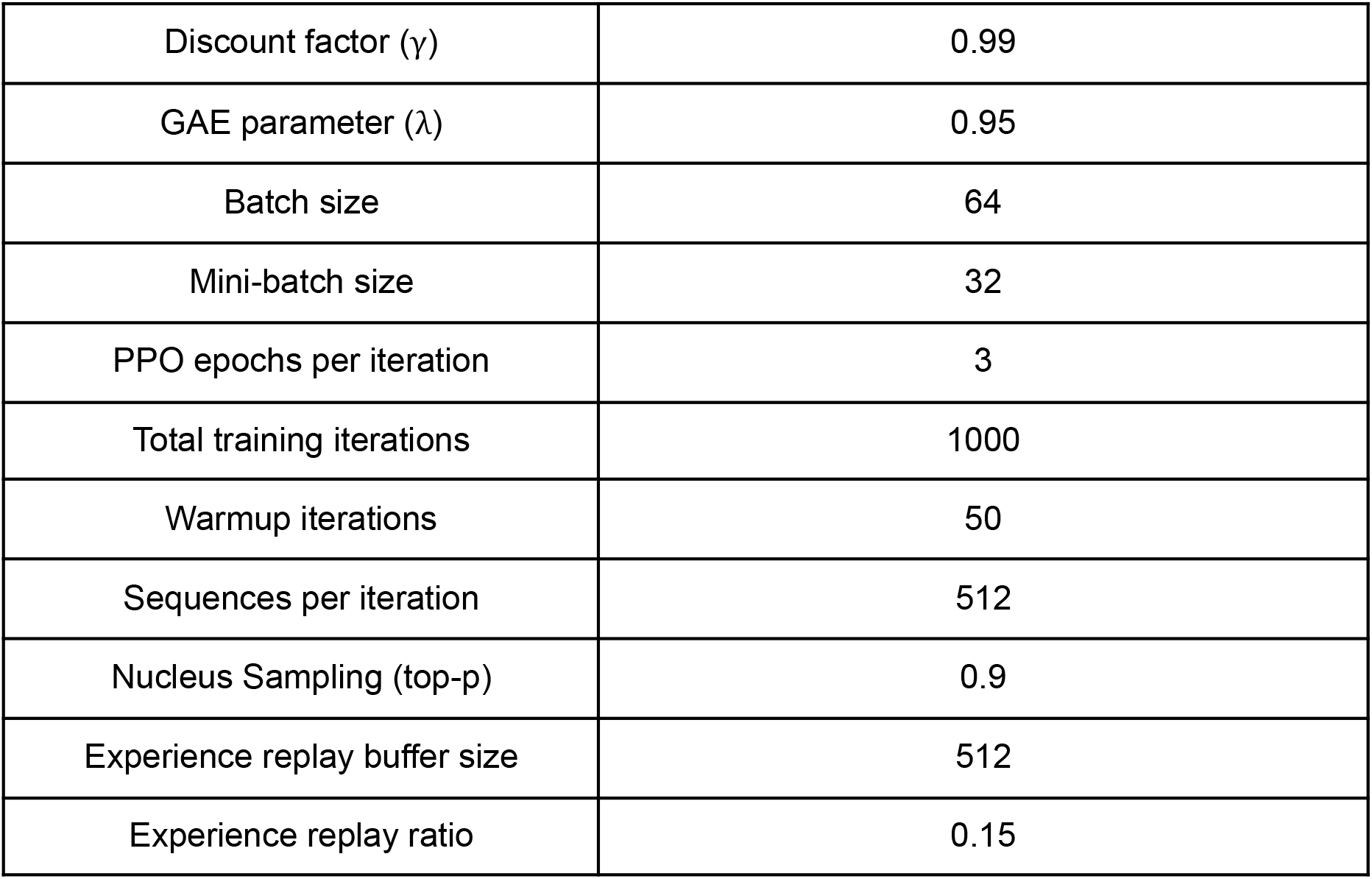
Detailed RL training hyperparameters.

| Hyperparameter | Value |
| --- | --- |
| Policy learning rate | 5.00E-05 |
| Value learning rate | 8.00E-05 |
| Off-Target Penalty ($\lambda_{off}$) | 1.0 -> 5.0 (linear) |
| PPO clip ratio ($\epsilon$) | 0.2 |
| Value loss coefficient ($c_v$) | 0.5 |
| Entropy coefficient ($c_e$) | 0.05 |
| KL penalty coefficient ($c_{KL}$) | 0.01 |
| Target KL divergence | 0.04 |
| Discount factor ($\gamma$) | 0.99 |
| GAE parameter ($\lambda$) | 0.95 |
| Batch size | 64 |
| Mini-batch size | 32 |
| PPO epochs per iteration | 3 |
| Total training iterations | 1000 |
| Warmup iterations | 50 |
| Sequences per iteration | 512 |
| Nucleus Sampling (top-p) | 0.9 |
| Experience replay buffer size | 512 |
| Experience replay ratio | 0.15 |

**Supplement Video 1: See Github:**https://github.com/zeyuChenLab/GO-CRE.git.

